# A Cophylogenetic Approach for Virus-Host Interaction Prediction

**DOI:** 10.64898/2026.02.26.708038

**Authors:** Md Zarzees Uddin Shah Chowdhury, T. M. Murali, Palash Sashittal

## Abstract

Advances in metagenomics have rapidly expanded viral discovery, revealing vast diversity across Earth’s virosphere. Yet most virus–host interactions—i.e., which viruses infect which hosts—remain unrecorded. Identifying these interactions is essential for anticipating zoonotic spillover events and advancing biomedical applications such as bacteriophage therapy. However, the sheer diversity of viruses and hosts makes comprehensive experimental mapping infeasible, motivating the need for computational approaches. Most existing prediction methods rely on supervised learning strategies that use sequence-derived features, such as codon usage bias or *k*-mer frequencies, and do not model the coevolutionary processes that shape virus–host interactions. This limits their ability to generalize and the evolutionary interpretability of their predictions. We introduce CoEvoLink, a framework for predicting virus–host interactions that integrates sequence-based evidence with phylogenetic signal by explicitly modeling the coevolutionary histories of viruses and hosts. CoEvoLink infers likely but unobserved interactions by minimizing the number of evolutionary events required to explain them, yielding the most parsimonious interaction under a coevolutionary model. This formulation generalizes classical maximum parsimony, typically defined on a single phylogeny, by jointly optimizing parsimony across both virus and host phylogenies. Sequence-based information is incorporated by assigning a cost to each potential interaction that reflects its likelihood based on genomic features. By drawing a connection between computing parsimony on interaction matrices and maximum parsimony on phylogenetic networks, we derive a polynomial-time algorithm that balances parsimony with sequence-derived prediction cost. We demonstrate the effectiveness of CoEvoLink on simulated data under diverse coevolutionary models. Applying CoEvoLink, we identified putative bat hosts of betacoronaviruses that have not yet been cataloged in the VIRION database. On a benchmark derived from metagenomic sequencing data, we demonstrate that CoEvoLink improves the performance of existing phage-host prediction tools using cophylogenetic information.

**Code availability:** https://github.com/sashittal-group/CoEvoLink

Note: This paper is accepted at RECOMB 2026 (30th Annual International Conference on Research in Computational Molecular Biology).

## 1 Introduction

Recent advances in metagenomic sequencing have led to the rapid discovery of new viruses. However, current databases cataloging known virus-host interactions, i.e. which viruses can infect which hosts, are very sparse with most interactions likely unrecorded [1–3]. Resolving these unknown virus-host interactions is critical for public health and biomedical applications. For instance, uncovering the potential hosts of known and novel viruses is a crucial component of identifying potential zoonotic spillover events and preventing future pandemics [4–6]. Identifying the hosts of bacteriophages, i.e., viruses that infect bacteria, has important biomedical applications such as developing phage therapies to combat antibiotic-resistant infections [7–10]. The immense diversity of both viruses and potential hosts makes experimental identification of all possible viral-host interactions intractable. Hence, there is a need for computational methods that can predict likely interactions and help prioritize their experimental validation [11].

Virus-host interactions are shaped by the complex process of *coevolution*, where viruses and hosts induce reciprocal selective pressures and evolve in response to each other [12–14]. Researchers have developed several *cophylogenetic* models and methods to study the interconnected evolutionary histories of symbionts and hosts [15, 16], where phylogenetically similar viruses typically infect phylogenetically similar hosts. As such, the shared evolutionary history of viral host repertoires and host viromes provides a crucial signal to identify unknown virus-host interactions.

Most existing computational approaches for predicting virus-host interactions do not account for their co-evolutionary history [17–23]. Current methods rely largely on supervised learning strategies that analyze viral genomes and leverage sequence-based features, such as codon usage bias [24–26], nucleotide composition [27], or *k*-mer frequencies [28, 29] to predict potential hosts of viruses. These features are informative because viruses often adapt their genomes to resemble those of their hosts [26, 30, 31]. While these methods have shown utility, they suffer from several limitations. First, they typically require viral sequences as input and can only test against hosts present in pre-defined reference databases, limiting their generalizability [32]. Second, they often demand substantial computational resources and expertise to retrain models whenever new host data are added. Most importantly, these approaches offer limited interpretability, i.e. they predict interactions without providing evolutionary insight into why a virus can infect a host. These limitations highlight the need for methods that can integrate the two complementary sources of information: genomic sequence-based features and phylogenetic signal from the co-evolutionary histories of viruses and hosts.

To address this need, we propose a novel framework to predict virus-host interactions that integrates sequence-based information with cophylogenetic signal by explicitly modeling their coevolutionary histories. We formulate the problem of inferring likely but currently unknown interactions between viruses and hosts while minimizing the number of evolutionary events needed to explain them, thereby yielding the most parsimonious interactions under a given coevolutionary model. Our formulation generalizes the traditional notion of maximum parsimony which is usually defined on a single phylogeny by maximizing parsimony across the host and the virus phylogenies simultaneously. We incorporate sequence-based information by assigning a cost to each potential interaction which reflects the likelihood inferred from genomic features — higher cost indicating lower sequence-based support. Our approach yields a natural trade-off between the number of predicted interactions (or the total cost of the predictions) and parsimony score of the interactions. We derive a polynomial time algorithm to balance this trade-off by drawing connections to a maximum parsimony problem on phylogenetic networks. The resulting method, CoEvoLink, is computationally efficient, interpretable, and readily integrable with existing sequence-based approaches.

We demonstrate the effective performance of CoEvoLink on simulated and real data. On simulated data with different coevolutionary models we show that CoEvoLink can recover missing virus-host interactions even under substantial sparsity in known interaction while relying only on cophylogenetic signal. We applied Co-EvoLink to uncover putative bat hosts of betacoronaviruses that are missing in the VIRION database [33]. CoEvoLink predictions recapitulate known bat-betacoronavirus associations and identified 10 bat hosts that may be reservoirs for betacoronaviruses from subgenera that contain viruses that infect humans, exhibiting spillover potential. Lastly, we evaluate the performance of CoEvoLink in identifying potential hosts for bacteriophages derived from metagenomic sequencing data. We show that CoEvoLink leverages coevolutionary signal to improve the performance of existing phage-host interaction prediction tools.

## 2 A Cophylogenetic Approach to Predict Virus-Host Interactions

We represent the virus-host interactions between *n* viruses and *m* hosts by a *n × m* interaction matrix *A*, where *a* _*i,j*_ = 1 if virus *i* and host *j* interact, i.e. virus *i* can infect host *j*, and *a* _*i,j*_ = 0 otherwise (Fig. 1). The *host range* of a virus *j* is the set of hosts it can infect (*a* _*i,j*_ = 1) and the *virome* of a host *i* is the set of viruses *j* that are capable of infecting it (*a* _*i,j*_ = 1). We represent the evolutionary relationships among viruses and among hosts by their respective phylogenies. Specifically, the viral tree *T*_*v*_ is a rooted phylogenetic tree, in which each internal vertex represents an ancestral virus and the leaf set *L*(*T*_*v*_) represent *n* extant viruses that correspond to the rows of *A*. Similarly, the host tree *T*_*h*_ is a rooted phylogenetic tree describing the evolution of the *m* hosts corresponding to the columns of *A*.

**Fig. 1.**
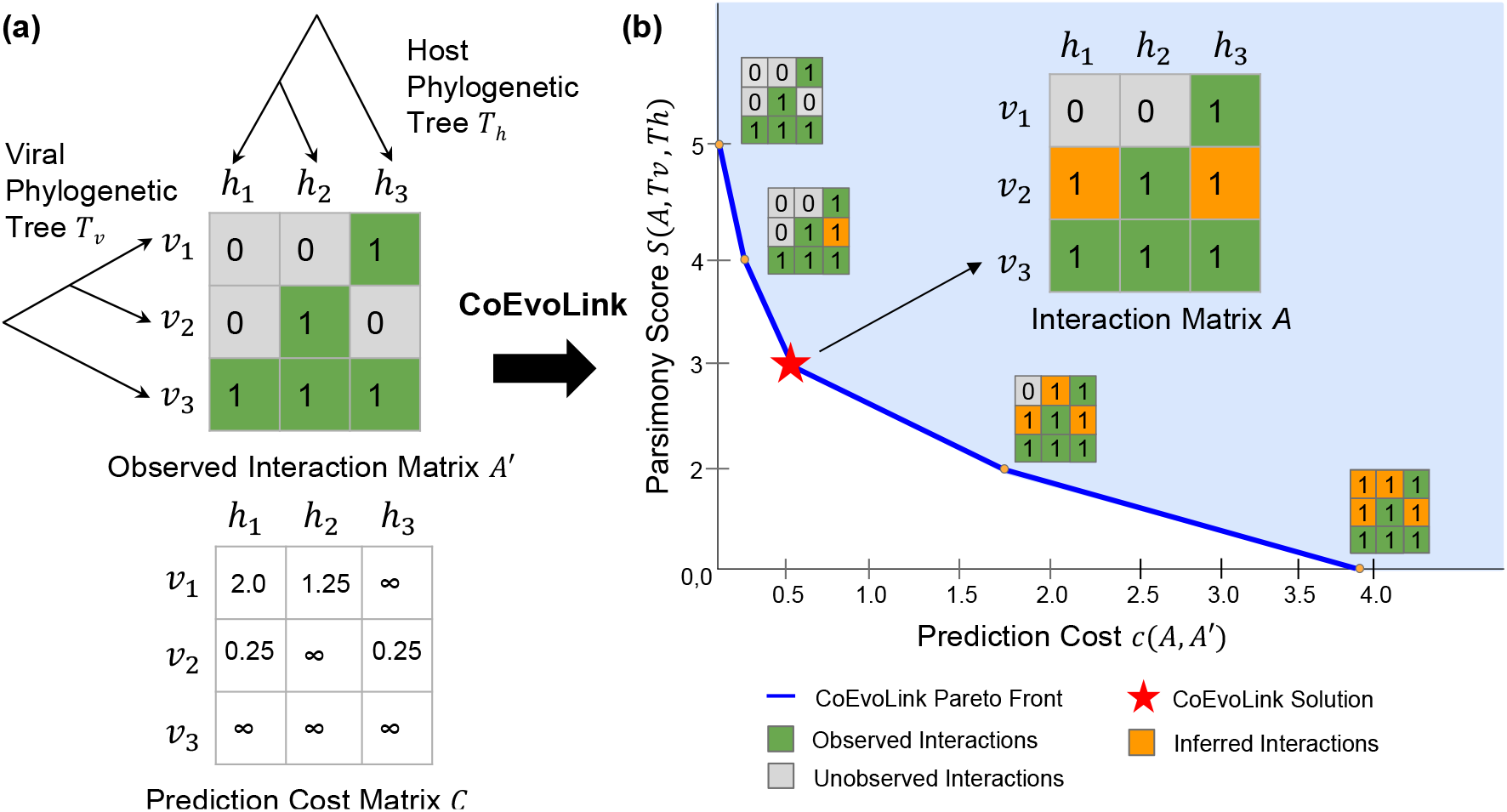
Virus-host interaction prediction using CoEvoLink. (a) CoEvoLink takes an observed binary interaction matrix *A*′ as input, where 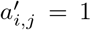 indicates a documented virus–host interaction and 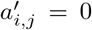 = 0 denotes either an untested interaction or a true non-interaction. The method infers the true interaction matrix A by proposing 0 → 1 flips in *A*′, guided by the viral and host phylogenies (*T*_*v*_ and *T*_*h*_) and a prediction cost matrix *C*, where *c* _*i,j*_ reflects the likelihood that virus *i* interacts with host *j*. (b) CoEvoLink reveals the Pareto front by balancing parsimony on the phylogenies, *S*(*A, T*_*v*_, *T*_*h*_), against the prediction cost *c*(*A, A′*). The most plausible interaction matrix is identified by computing the *elbow* of the Pareto front.

The interactions between only a fraction of all possible host-virus pairs have been studied. Hence, the known interaction matrix *A* ^*′*^ is a very sparsely observed version of the true interaction matrix *A*, where the documented interactions have strong evidence, i.e. we observe 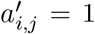 only if it is a true interaction *a* _*i,j*_ = 1 (Fig. 1a). However, the interaction of several compatible host-virus pairs has not been tested, and as such, several host-virus pairs (*i, j*) with 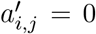 represent unknown true interactions (*a* _*i,j*_ = 1). Our goal is to identify such host-virus pairs, i.e., perform 0 → 1 flips in the observed interaction matrix *A* ^*′*^, to recover the true interaction matrix *A*.

The interaction matrix *A* between the extant viruses and hosts is the result of a long history of coevolution. We represent the history of virus-host interactions by the *extended interaction matrix* 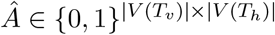, where each entry 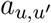 represents if virus *u* ∈ *V* (*T*_*v*_) (which may be ancestral or extant) can infect host *u* ^*′*^ ∈ *V* (*T*_*h*_) (also ancestral or extant). Importantly, the extended interaction matrix *Â* must be *consistent* with the interaction matrix *A*, i.e., *â* _*i,j*_ = *a* _*i,j*_ for each extant host *i* ∈ *L*(*T*_*h*_) and extant virus *j* ∈ *L*(*T*_*v*_). Since multiple extended interaction matrices may be consistent with a given interaction matrix, we require an evolutionary model to evaluate and compare the alternative extensions.

Several models developed to study evolution of parasite-host interactions focus on congruence between parasite and host phylogenies [34–37]. Specifically, reconciliation-based models reconstruct the co-evolutionary history of parasites and hosts by inferring a mapping from vertices of the phylogeny of parasites to vertices of phylogeny of hosts [38]. However, these models assume that each parasite can only interact with a unique host. As such, these models are not applicable for inferring putative virus-host interactions since an increasing body of evidence suggests that several viruses have wide host ranges [39–41].

An alternative approach that allows a virus to have multiple hosts is to treat the virus-host interactions as phylogenetic characters [16, 42–44]. Similar to Braga et al. [16], we treat the membership of each virus *i* in the virome of the hosts as a two-state phylogenetic character along the host tree with states: 0 and 1. Similarly, we treat the membership of a host in the host range of a virus as a two-state phylogenetic character along the virus tree. Evolutionary changes in the viruses and hosts can lead to gain (0 → 1 changes) or loss (1 → 0 changes) of virus-host interactions along the two phylogenies.

We take the maximum parsimony approach [45] to quantify the cophylogenetic signal of an interaction matrix, under the assumption that mutations that cause changes in host-virus interactions are rare. Specifically, we define the parsimony score of an interaction matrix as the minimum number of evolutionary changes in virus-host interactions along the virus and the host phylogenies that explain the observed interactions at the leaves. Formally, we define the parsimony score of an extended matrix *Â* with respect to the viral tree *T*_*v*_ as

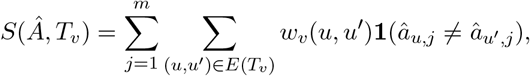

where **1** is the indicator function and *w* _*v*_(*u, u* ^*′*^) is the non-negative weight associated with edge (*u, u* ^*′*^) in the viral tree *T*_*v*_ depicting the likelihood for a state transition to occur along edge (*u, u* ^*′*^). Thus, *S*(*Â, T* _*v*_) is obtained by summing, over all hosts *j*, the total weight of all edges (*u, u* ^*′*^) in *T*_*v*_ where host *j* interacts with *u* but not with *u* ^*′*^ or *vice versa*.

Analogously, we consider the transpose *Â* ^*T*^ of the interaction matrix to define the parsimony score *S*(*Â* ^*T*^, *T*_*h*_) with respect to the host tree *T*_*h*_ with edge weights *w* _*h*_. The total parsimony score *S*(*A, T*_*v*_, *T*_*h*_) of an interaction matrix *A* with respect to the viral tree *T*_*v*_ and host tree *T*_*h*_ is

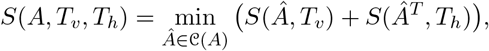

where 𝒞(*A*) is the set of consistent extended interaction matrices of *A*. This formulation extends classical maximum parsimony, in which a character matrix is evaluated on a single phylogenetic tree [46, 47], by jointly modeling the evolution of virus–host interaction states across two phylogenies and minimizing the total number of interaction gains and losses over both trees.

Besides the cophylogenetic signal, genomic sequence-based features are also informative of virus-host interactions. Several methods [24, 27, 28] have been developed to derive interaction prediction scores based on genomic features. We incorporate this information by associating each virus-host pair (*i, j*) with cost *c* _*i,j*_ derived from such scores, such that *c* _*i,j*_ is low if interaction of virus *i* and host *j* is strongly supported by genomic features and thus (*i, j*) is more likely to have an undocumented interaction. We denote the total cost of all the predictions, i.e. 0 → 1 flips in the interaction matrix, by 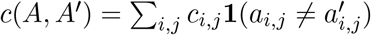.

There exists a tradeoff between the prediction cost and the parsimony of an interaction matrix. Observe that the parsimony score can always be decreased as more 0 →1 flips are allowed in the interaction matrix. While the lowest prediction cost (*c*(*A, A* ^*′*^) = 0) is attained when we do not perform any 0→ 1 flips, we achieve the lowest possible parsimony score *S*(*A, T*_*v*_, *T*_*h*_) = 0 when all 0 entries in the matrix are flipped to 1 (Fig. 1b). We must balance this trade-off to control the inference of false-positive host-virus interactions.

One approach to balance the trade-off between two objectives is the weighted sum approach [48]. In this approach, we minimize a convex combination of the parsimony score *S*(*A, T*_*h*_, *T*_*v*_) and the total cost *c*(*A, A* ^*′*^) of the 0→ 1 flips, where their relative contributions are determined by a parameter *λ* ∈ [0, 1]. This motivates the following Balanced Parsimonious Interaction Prediction Problem.

### Problem 2.1

(Balanced Parsimonious Interaction Prediction (B alanced PIP)). *Given an observed interaction matrix A* ^*′*^, *host tree T* _*h*_ *with edge weights w* _*h*_, *virus tree T* _*v*_ *with edge weights w* _*v*_ *and pa-rameter λ* ∈ [0, 1], *perform* 0 → 1 *flips in A* ^*′*^ *such that the resulting interaction matrix A minimizes λS*(*A, T*_*v*_, *T*_*h*_) + (1 − *λ*)*c*(*A, A* ^*′*^).

An alternative approach to balance the trade-off between *S*(*A, T*_*v*_, *T*_*h*_) and *c*(*A, A* ^*′*^) is to minimize the *S*(*A, T*_*v*_, *T*_*h*_) while introducing a parameter *δ* that bounds *c*(*A, A* ^*′*^) from above, i.e., *c*(*A, A* ^*′*^) ≤ *δ*. We formally pose the Bounded Parsimonious Flipping Problem as follows.

### Problem 2.2

(Bounded Parsimonious Interaction Prediction (B ounded PIP)). *Given an observed interac-tion matrix A* ^*′*^, *host tree T* _*h*_, *virus tree T* _*v*_ *and parameter δ* ∈ ℝ, *perform* 0 → 1 *flips in A* ^*′*^ *such that the resulting interaction matrix A minimizes S*(*A, T*_*v*_, *T*_*h*_) *and c*(*A, A* ^*′*^) ≤ *δ*.

We say that an inferred interaction matrix *A* is *Pareto optimal* [49] if it is better than all other interaction matrices in at least one of the two objectives, *S*(*A, T*_*v*_, *T*_*h*_) and *c*(*A, A* ^*′*^). In other words, there is no other interaction matrix that has both lower parsimony score and lower cost of predictions. We pose the problem of inferring all Pareto optimal interaction matrices as follows.

### Problem 2.3

(Pareto Optimal Interaction Prediction (POIP)). *Given an observed interaction matrix A* ^*′*^, *host tree T* _*h*_, *virus tree T* _*v*_, *find all interaction matrices A such that (i) A can be obtained from A* ^*′*^ *by performing only* 0 → 1 *flips, (ii) A is Pareto optimal with respect to S*(*A, T*_*v*_, *T*_*h*_) *and c*(*A, A* ^*′*^).

## 3 Characterizing Parsimonious Virus-Host Interactions

We develop the theoretical foundations for computing maximum parsimony of interaction matrices defined over two phylogenies. Unlike classical maximum parsimony formulations that operate on a single phylogeny, our framework involves maximizing total parsimony of the interaction matrix across two phylogenetic trees — one for viruses and one for hosts. We establish formal connections between this framework and maximum parsimony on phylogenetic networks. Moreover, we show that the two alternative ways of balancing the trade-off between the prediction cost and parsimony of an interaction matrix (Prob. 2.1 and Prob. 2.2), differ in computational complexity. Specifically, we demonstrate that the Balanced PIP problem (Prob. 2.1) can be reduced to the Minimum *s*–*t* Cut problem and therefore admits a polynomial-time algorithm, whereas the Bounded PIP problem (Prob. 2.2) is NP-hard.

Phylogenetic networks are graphs that are used to describe non tree-like evolutionary relationships in phylogenetics, such as those arising from recombination or horizontal transfer [50, 51]. Formally, a phylogenetic network is a connected weighted directed acyclic graph where each leaf (vertex with 0 outgoing edges) is labeled by 0 or 1. Computing the maximum parsimony score of a network involves labeling the internal vertices with 0 or 1 such that total weight of all edges where the labels of the source and target vertices differ is minimized. In the literature, this quantity is referred to as *hardwired parsimony score* [52] of the network. We will show that the virus tree *T*_*v*_ and the host tree *T*_*h*_ can combined into a single phylogenetic network *N* (*A, T*_*v*_, *T*_*h*_) with leaves labeled by entries of *A* such that computing the parsimony *S*(*A, T*_*v*_, *T*_*h*_) of the interaction matrix is equivalent to computing the hardwired parsimony of *N* (*A, T*_*v*_, *T*_*h*_).

Intuitively, *N* (*A, T*_*v*_, *T*_*h*_) contains *m* copies of the virus tree and *n* copies of the host tree, where leaves corresponding to the same virus-host pair (*i, j*) are merged into a single vertex and labeled by *a* _*i,j*_ (Fig. 2). Formally, we construct *N* (*A, T*_*v*_, *T*_*h*_) as follows. We introduce a vertex (*x, i*) for each host *i* and internal vertex *x* of the virus tree, a vertex (*j, y*) for each virus *j* and internal vertex *y* of the host tree, and a leaf (*i, j*) for each virus *i* and host *j* labeled by *a* _*i,j*_. We introduce an edge ((*x, i*), (*x* ^*′*^, *i*)) with weight *w* _*v*_(*x, x* ^*′*^) for each edge (*x, x* ^*′*^) in virus tree *T*_*v*_ and host *i* and an edge ((*j, y*), (*j, y* ^*′*^)) with weight *w* _*h*_(*y, y* ^*′*^) for each edge (*y, y* ^*′*^) in host tree *T*_*h*_ and virus *j*. Lastly to make the network rooted, we introduce root vertex *r* and add weight 0 edges from *r* to vertices (*r*(*T*_*v*_), *i*) for each virus *i* and (*j, r*(*T*_*h*_)) for each host *j*. This results in a phylogenetic network with 1 + *m* |*V* (*T*_*v*_) | + *n*|*V* (*T*_*h*_)| − *nm* vertices and *m* |*E*(*T*_*v*_)| + *n*|*E*(*T*_*h*_)| + *n* + *m* edges that can be constructed in time linear in the size of the interaction matrix. *N* (*A, T*_*v*_, *T*_*h*_) is related to parsimony score of interaction matrix as follows.

**Fig. 2.**
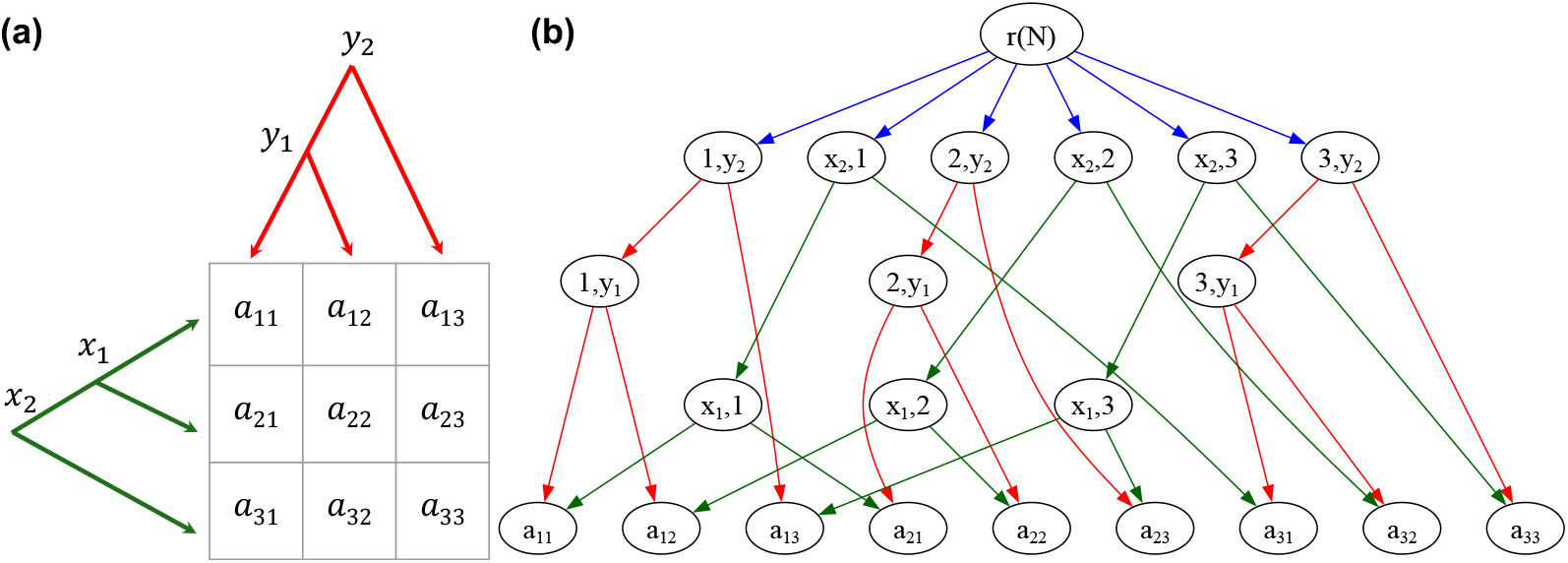
Example showing construction of phylogenetic network *N* (*A, T*_*v*_, *T*_*h*_). (a) A *n × m* Interaction matrix *A* with viral phylogeny *T*_*v*_ (with *n* leaves corresponding to the *n* rows of *A*) and host phylogeny *T*_*h*_ (with *m* leaves corresponding to the *m* columns of *A*). (b) Phylogenetic network *N* (*A, T*_*v*_, *T*_*h*_) with 1 + *m*|*V* (*T*_*v*_)| + *n*|*V* (*T*_*h*_)| − *nm* vertices and *m*|*E*(*T*_*v*_)| + *n*|*E*(*T*_*h*_)| + *n* + *m* edges.

**Fig. 3.**
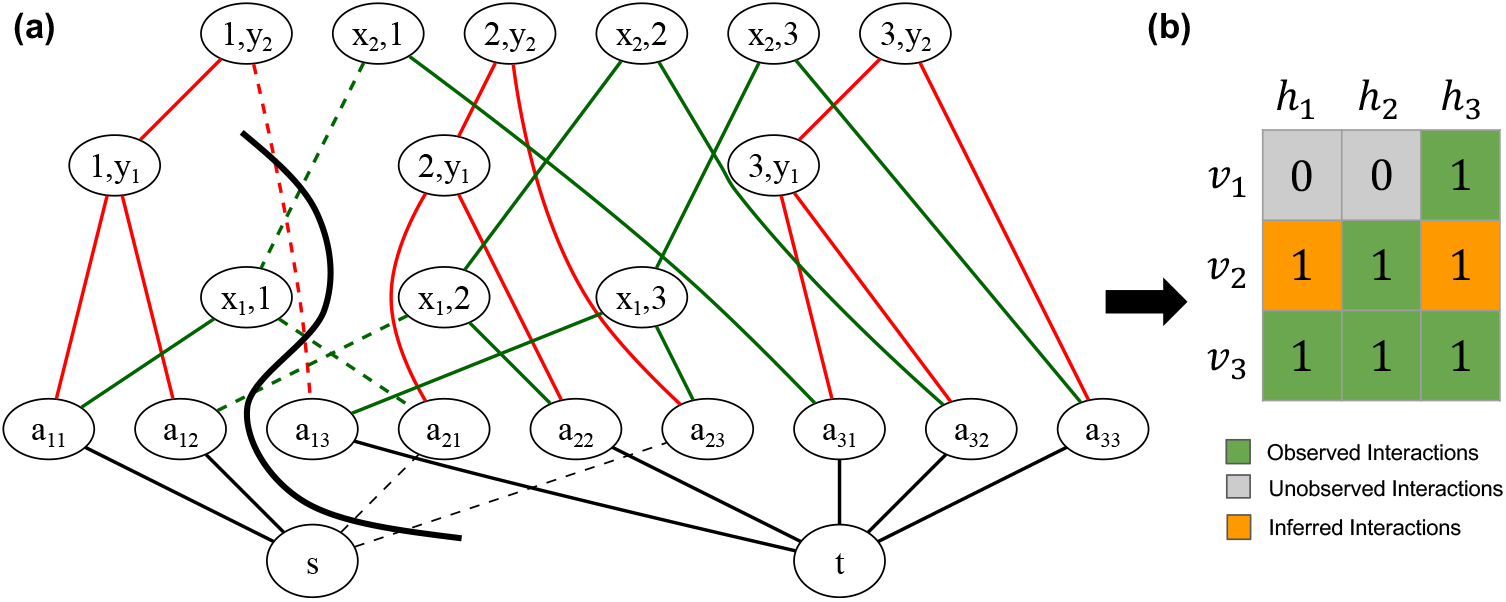
Schematic showing the correspondence between *s*-*t* cut of a graph *G* and predicted virus-host interaction matrix. (a) *G*(*A* ^*′*^, *T*_*v*_, *T*_*h*_) for interaction matrix *A* ^*′*^ shown in Fig. 1. An *s*-*t* cut of *G*(*A* ^*′*^, *T*_*v*_, *T*_*h*_) separates the graph into two components which determines the values of the entries in the (b) inferred interaction matrix. Entries *a* _1,1_ and *a* _1,2_ belong to the component with vertex *s* and are thus labeled by 0 and the rest of the entries are labeled by 1.

### Lemma 3.1.

*The parsimony score S*(*A, T*_*v*_, *T*_*h*_) *of an interaction matrix A is equal to the hardwired parsimony score of phylogenetic network N* (*A, T*_*v*_, *T*_*h*_).

Maximum parsimony is a widely studied problem on both phylogenetic trees and networks. The maximum parsimony score for a tree can be computed in polynomial time using the well-known Fitch’s algorithm [47] or Sankoff’s algorithm [46]. While it was shown that extending these algorithms does not compute the accurate parsimony score for phylogenetic networks in general [53], the specific structure of *N* (*A, T*_*v*_, *T*_*h*_) where every vertex with more than one incoming edge (*reticulation vertex*) is a leaf, allows us to extend Sankoff’s algorithm to compute the parsimony score of *N* (*A, T*_*v*_, *T*_*h*_) in *O*(*nm*) time. Due to the equivalence described above (Lemma 3.1), we have the following corollary (Proof in Supp. Sec. A).

### Corollary 3.1.

*The parsimony score S*(*A, T*_*v*_, *T*_*h*_) *of an interaction matrix A with n viruses and m hosts can be computed in O*(*nm*) *time*.

Leveraging the connection to maximum parsimony on phylogenetic networks, we show that the Balanced Parsimonious Interaction Prediction Problem (Prob. 2.1) can be reduced to the well-known minimum *s*-*t* cut problem. Given a weighted undirected graph *G* with vertices *s* and *t*, the minimum *s*-*t* cut problem is to find a subset of edges (the *cut*) with minimum total weight whose removal results in a graph with *s* and *t* in separate components. The minimum *s*-*t* cut problem can be solved in *O*(|*V* (*G*) ||*E*(*G*) |) time using a flow-based algorithm [54] that relies on the max-flow min-cut theorem [55].

We build a graph *G*(*A* ^*′*^, *T*_*v*_, *T*_*h*_) for an observed interaction matrix *A* ^*′*^ in which the two components induced by an *s*-*t* cut determine the entries of inferred interaction matrix *A* and the size of the cut corresponds to the convex combination *λS*(*A, T*_*v*_, *T*_*h*_) + (1 − *λ*)*c*(*A, A* ^*′*^) of parsimony and prediction cost (see Fig. for an example). *G*(*A* ^*′*^, *T*_*v*_, *T*_*h*_) is obtained by modifying network *N* (*A* ^*′*^, *T*_*v*_, *T*_*h*_) with the following four steps. First, the root node and orientations of edges of the network are removed. Second, all edge weights are scaled by *λ*. Third, source vertex *s* is added and connected to every vertex (*i, j*) where 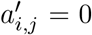 with edge weight (1 − *λ*)*c* _*i,j*_. Fourth, target vertex *t* is added and connected to every vertex (*i, j*) where 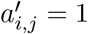 with edge weight ∞. We show that the minimum *s*-*t* cut in *G*(*A* ^*′*^, *T*_*v*_, *T*_*h*_) provides the solution to the Balanced PIP problem (proof in Supp. Sec. A.2), which leads to the following result.

### Theorem 3.1.

*The Balanced Parsimonious Interaction Prediction Problem can be solved in O*(*n* ^2^*m* ^2^) *time, where n is the number of viruses and m is the number of hosts*.

We also show that the Bounded Parsimonious Interaction Problem (Prob. 2.2) is NP-hard, even when for the cases where *n* = 1 or *m* = 1, using a reduction from the Knapsack problem A.3.

### Theorem 3.2.

*The Bounded Parsimonious Interaction Problem is NP-hard, even for n* = 1 *or m* = 1. We provide proofs of Theorem 3.1 and Theorem 3.2 in the Supplement (Sec. A).

## 4 CoEvoLink: Flow-based Algorithms for Virus-Host Interaction Prediction

We introduce CoEvoLink, a suite of novel algorithms based on the flow-based method for solving the Balanced PIP problem (Theorem 3.1) to predict putative virus-host interactions while incorporating phylogenetic and sequence-based information. CoEvoLink has four modules: (1) enumerating Pareto optimal interaction matrices, (2) determining the number of missing interactions by balancing the trade-off between parsimony and prediction cost, (3) solving the B ounded PIP problem, i.e. computing the most parsimonious interaction matrix with prediction cost below a given threshold *δ*, and (4) Performing taxonomy-level predictions. We describe module (4) here, while modules (1), (2) and (3) are detailed in Supp. Sec. B

### Performing taxonomy-level predictions

In many applications, it is useful to predict virus-host interactions not only at the species level but also across high taxonomic ranks – for example, whether a virus is likely to infect any member of a particular genus or family. Since CoEvoLink explicitly models the co-evolution of virus-host interactions, it is able to make taxonomy-level predictions. Specifically, CoEvoLink predicts virus *i* infects a host taxon if the most recent common ancestor *u* of hosts from that taxon in the host phylogeny interacts with virus *i*, i.e. *Â* _*i,u*_ = 1 in the extended interaction matrix *Â*. Intuitively, CoEvoLink predicts a virus-taxon interaction when a sufficient number of hosts within that taxon are predicted to interact with the virus under the most parsimonious evolutionary scenario. This approach allows predictions at any taxonomic level by aggregating over phylogenetically similar hosts.

## 5 Results

### 5.1 Simulated data

We performed simulations to evaluate the ability of CoEvoLink to recover virus-host interactions using only co-phylogenetic signal, without sequence-based information. Our simulations are based on phylogenies of *n* = 94 coronaviruses (family Coronaviridae) and *m* = 233 bats (order Chiroptera). We obtained the virus and bat genomes from NCBI and constructed phylogenies using IQ-TREE [56] (details in Supplement E and S7a Figure S7b). We generated two sets of simulations using distinct evolutionary models for virus-host interactions. In the first set, virus-host interactions evolve under the Mk model [57] with two-state characters (0 = no interaction and 1 = interaction). We varied state transition rates *ρ* ∈ {0.1, 0.2, 0.3, 0.4, 0.5} to represent different evolutionary dynamics and introduced missing interactions by randomly performing 1 → 0 flips for *d* = {10%, 20%, 50%} of true interactions. In the second set of simulations, we employed the *host-repertoire evolution* model described in Braga et al. [16], where host colonization occurs with a three-state model: 0 = no interaction, 1 represents that virus has the ability to interact with a host, and 2 represents virus-host interaction. We present the results on Mk-model simulations here and describe the results on the host-repertoire model in Supplementary Sec. C.2.

We evaluate CoEvoLink under two settings. In the *unsupervised* setting, the number of missing true interactions is unknown; CoEvoLink jointly estimates this number and recovers the interaction matrix by selecting the elbow of the Pareto front—avoiding manual thresholding required by existing approaches. In the *supervised* setting, the true number of missing interactions is provided, and CoEvoLink returns the most parsimonious interaction matrix consistent with that number of predictions.

Our simulations demonstrate that CoEvoLink effectively recovers missing virus–host interactions by exploiting cophylogenetic signal. In the unsupervised setting, CoEvoLink achieves a median F1-score of 0.633 (Fig. 4a), driven by near-perfect recall (0.946; Fig. S2b) and moderate precision (0.52; Fig. S2a). This pattern reflects a conservative strategy in which the method prioritizes parsimony and therefore tends to predict more interactions, resulting in lower precision when the true missing rate is small (Supplementary Fig. S2a). The number of predicted interactions can be tuned by modifying how the *elbow* is defined and detected on the Pareto front (Fig. 4c), providing a principled way to balance false positives and false negatives. When the true number of missing interactions is known, performance improves substantially: CoEvoLink reaches a median F1-score of 0.772 across all simulation conditions (Fig. 4b). We observe similar trends under simulations generated with the three-state *host-repertoire model* (Supplementary Sec. C.2), indicating that the method generalizes across varying interaction-evolution models. These results highlight that CoEvoLink reliably infers missing virus–host interactions by capturing the underlying cophylogenetic structure.

**Fig. 4.**
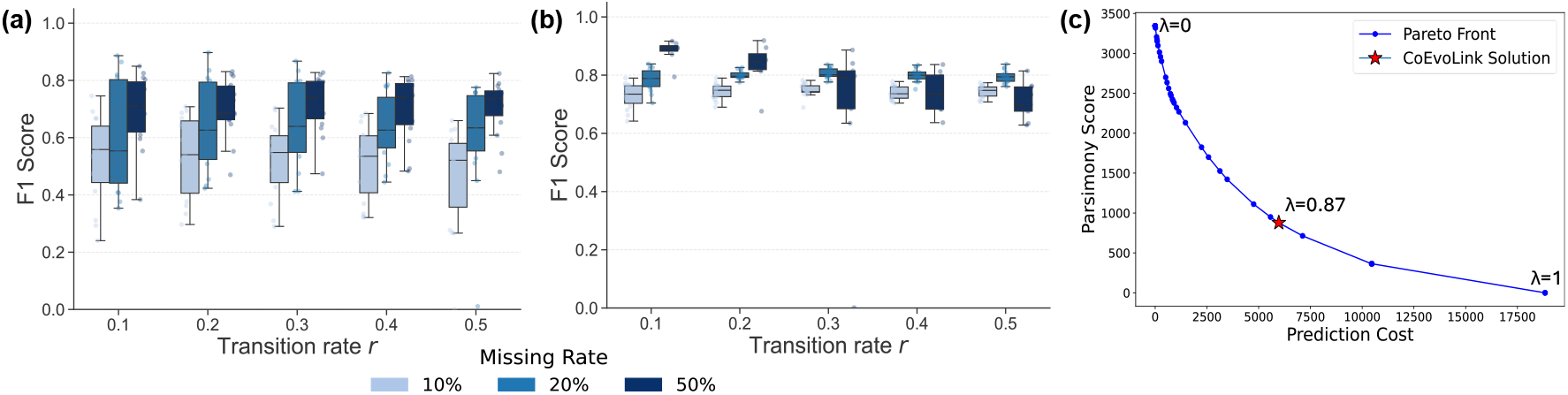
CoEvoLink identifies missing interactions in simulated data using cophylogenetic information. F1-score across different missing rates when number of missing interaction is (a) unknown (unsupervised) and (b) known (supervised). (c) Pareto front for one of the simulated instances identified by CoEvoLink.

### 5.2 CoEvoLink predicts putative coronavirus-bat interactions

We applied CoEvoLink to uncover putative bat hosts of betacoronaviruses that are missing in The Global Virome in One Network (VIRION) database [33]. This database is the largest open-access atlas of virus-vertebrate interactions, containing approximately 6% of total known vertebrate diversity and covering multiple viral families. In contrast, *β*CoV [58] is a more specialized dataset that catalogs genus-level annotations of 1116 bat species, indicating whether each species is recognized in the literature as a betacoronavirus host. VIRION contains 185 of these 1116 bat species. Of these, *β*CoV annotates 61 as betacoronavirus hosts. However, VIRION lacks any recorded interaction for 24 of these 61 bats with the 21 betacoronaviruses listed in that database, revealing clear gaps in the VIRION interaction network.

We evaluated CoEvoLink’s ability to make genus-level predictions on the 185 bats present in both VIRION and *β*CoV, using *β*CoV as the ground-truth. We downloaded genomes of these bats and the 21 betacoronaviruses in VIRION from NCBI and generated the phylogenies for both viruses (*T*_*v*_) and bats (*T*_*h*_) using IQ-Tree [56] (details in Supp. Sec. E). We deemed a bat species *j* to be a predicted betacoronavirus host if CoEvoLink inferred an interaction at the taxonomic rank of betacoronaviruses, i.e. if *ar*(*T*_*v*_),_*j*_ = 1.

We benchmarked CoEvoLink’s performance against an ensemble of eight predictive models developed by Becker et al. [58]. The ensemble integrates the prediction of three trait-based models (Trait-1, Trait-2, Trait-3) that use ecological and physiological characteristics to predict potential hosts; four network-based models (Network-1, Network-2, Network-3, Network-4) that estimate link probabilities from the observed host–virus network; one hybrid model (Hybrid-1), which employs a two-step kernel ridge regression combining trait and phylogenetic information. This ensemble model was originally proposed to prioritize surveillance targets for undiscovered betacoronaviruses in bats [58] and serves as a comprehensive benchmark for evaluating CoEvoLink.

CoEvoLink identified bat species that are betacoronavirus hosts more accurately than the ensemble [58]. Using the prediction scores of the ensemble model to define the cost matrix, we applied CoEvoLink to identify Pareto-optimal interaction matrices (Supp. Fig. S5a). CoEvoLink achieved a higher area under precision-recall curve (AUPRC) compared to the ensemble (0.369 vs 0.278, Fig. 5a), with similar trends observed when compared to each of the individual models (Supp. Fig. S6). Furthermore, CoEvoLink outperformed two trait-based (trait-1 and trait-3) and all four network-based models using prediction cost matrices derived from the corresponding model’s prediction scores (Supp. Fig. S6). Notably, for the Hybrid and the four network-based models, using CoEvoLink led to more accurate predictions compared to the ensemble of all eight models (Supp. Fig. S5(e-i)). These results demonstrate that integration of cophylogenetic information with trait- or network-based prediction scores using CoEvoLink leads to more accurate predictions.

**Fig. 5.**
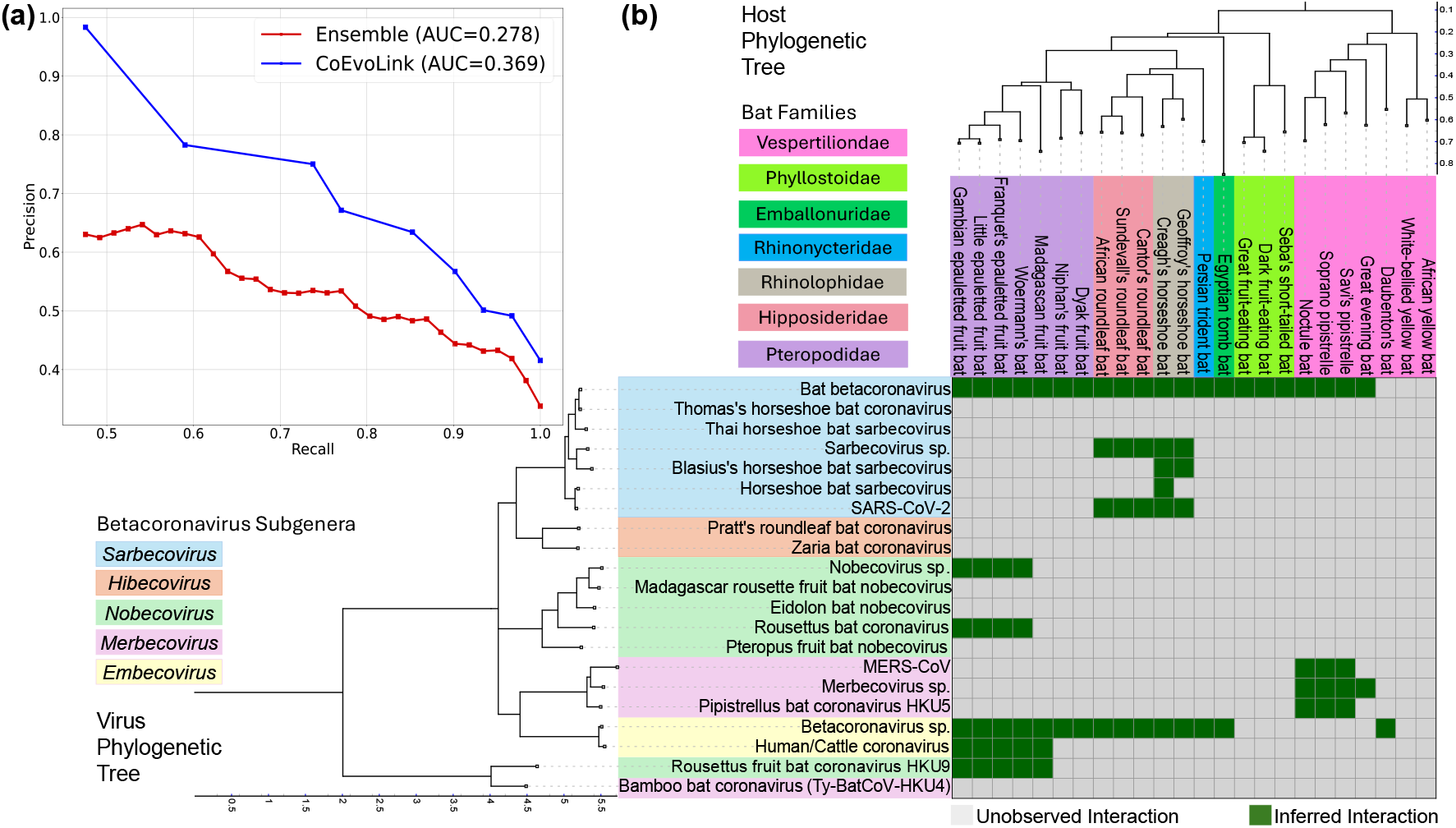
CoEvoLink outperforms existing methods in identifying bat species that are betacoronavirus hosts. (a) Precision-recall curve for CoEvoLink and the ensemble model on 185 bat species present in both VIRION and *β*CoV database. (b) Bat-betacoronavirus interactions predicted by CoEvoLink that are missing in the VIRION database.

We examined the species-level interactions inferred by CoEvoLink between betacoronaviruses and bats that are annotated as betacoronavirus hosts in *β*CoV but were missing from the VIRION database. CoEvoLink interaction matrix contained 77 interactions in total, including 34 involving *Sarbecoviruses*, 13 with *Nobecoviruses*, 10 with *Merbecoviruses*, and 20 with *Embecoviruses* (Fig. 5b). These predictions recapitulate known associations of bat families with betacoronavirus subgenera— Rhinolophidae are known hosts of *Sarbecoviruses*, Pteropodidae of *Nobecoviruses*, and Vespertilionidae of *Merbecoviruses*, respectively [59, 60] (Supp. Fig. S3). CoEvoLink identifies Bat betacoronavirus (VirusTaxID 1805463, *Sarbecovirus*) and Beta-coronavirus sp. (VirusTaxID 1928434, *Embecovirus*) as generalists with hosts spanning seven and five bat families, respectively. It also infers interactions between Hipposideridae and *Sarbecoviruses*—an association not typically emphasized but consistent with reports of SARS-like viruses persisting in Hipposideridae colonies and potentially contributing to spillover chains [61–65]. Notably, predicted interactions involving human-infecting betacoronaviruses (SARS-CoV-2 and Human/Cattle coronavirus OC43) highlight 10 bat species that may serve as overlooked reservoirs or intermediate hosts (Supp. Tables S1, S2). These predictions offer data-driven targets for ecological sampling and experimental assays to map the broader zoonotic landscape of betacoronaviruses.

### 5.3 CoEvoLink identifies potential hosts for novel bacteriophages

We evaluate the ability of CoEvoLink to predict hosts for bacteriophages using a benchmark derived from a MetaHiC sequencing dataset [66]. MetaHiC [67] is a metagenomic technique that provides evidence of virus-host interactions by capturing physical DNA-DNA associations between viruses and their bacterial hosts in mixed microbial populations. The benchmark comprises of 251 virus-host interactions collected from three distinct environments: 84 from human gut samples [67], 66 from cow fecal samples [68] and 101 from wastewater samples [69]. We employed IQTREE [56] to generate viral tree from the viral contigs and used taxonomy based host tree from bacteria derived from the MetaHiC data (details in Supp. Sec. E).

We compare CoEvoLink with three representative genomic-feature–based tools—PB-LKS [70], PHIST [71], and WIsH [72]. For each method, we obtained baseline predictions and converted them into prediction costs, where higher cost indicates lower prediction likelihood. CoEvoLink integrates these costs with parsimony to identify the most likely host for each phage. To evaluate prediction ranking, we used *top-k accuracy*, the fraction of phages whose true host appears among the *top k* predicted hosts (with *top-*1 as the single best prediction and *top-*3 allowing up to three candidates).

Across all three datasets, integrating CoEvoLink consistently improved or maintained the accuracy of existing host-prediction tools (Fig. 6). PB-LKS showed the largest improvements, with top-1 accuracy gains across all datasets (0.05 vs 0.012 for human gut, 0.06 vs 0.015 for cow fecal and 0.06 vs 0.01 for wastewater). For PHIST, top-1 accuracy improved for the wastewater dataset (0.495 with CoEvoLink vs 0.45 without) and on the human gut (0.43 with CoEvoLink vs 0.42 without) but same on the cow fecal (0.60) datasets. While top-1 accuracy of WIsH remained unchanged with or without CoEvoLink, CoEvoLink consistently improved top-3 accuracy of WIsH in all three datasets – 0.56 vs 0.5 for human gut, 0.71 vs 0.652 for cow fecal and 0.79 vs 0.76 for wastewater. Similar trends in top-*k* accuracy observed for the three methods across varying values of *k* (Supp. Fig. S4). These results demonstrate that CoEvoLink provides a framework to improve the performance of existing phage-host prediction tools in metagenomic studies.

**Fig. 6.**
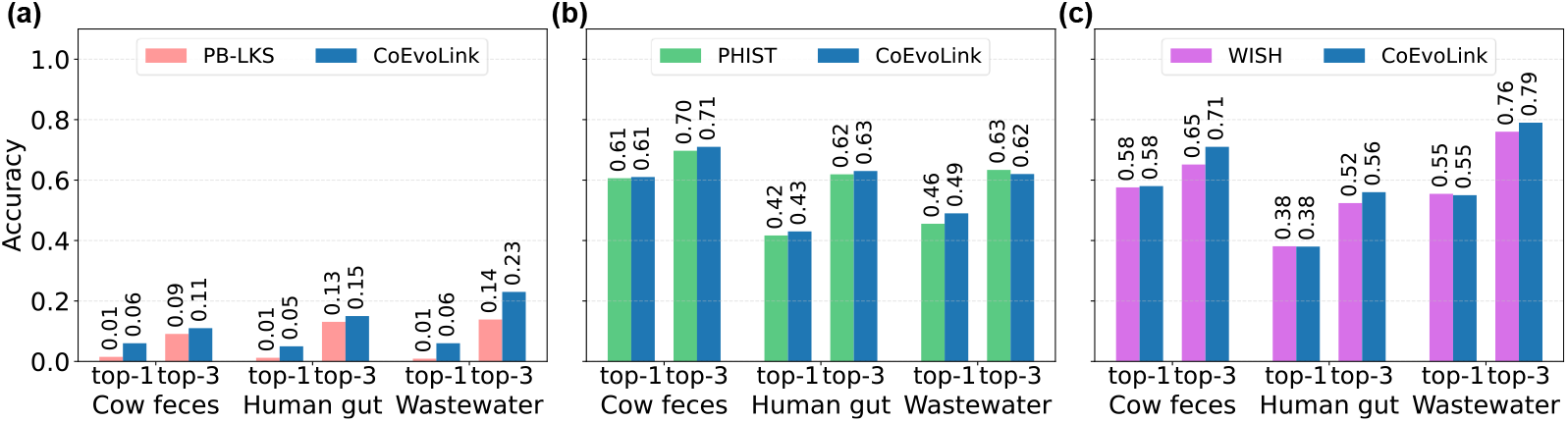
Integration with CoEvoLink improves the performance of sequence-based phage-host prediction methods. Accuracy of top-1 and top-3 predictions of (a) PB-LKS, (b) PHIST and (c)WIsH with and without CoEvoLink on predicting phage-host interactions derived from cow fecal, human gut and wastewater samples.

## 6 Discussion

We introduce CoEvoLink, a novel cophylogenetic framework to predict virus-host interactions. We formulate the problem of recovering missing virus-host interactions as finding an interaction matrix that minimizes the parsimony score, i.e. the number of evolutionary changes needed to explain the observed interactions, across both host and virus phylogenies. This maximum parsimony formulation generalizes traditional notion of maximum parsimony which is typically applied on a single phylogeny to jointly minimum parsimony across two phylogenies. CoEvoLink further balances this parsimony criterion with data-driven interaction costs, which can be derived from sequence similarity or from existing predictive models. This design enables CoEvoLink to incorporate cophylogenetic structure into any scoring-based prediction method. Importantly, the underlying B alanced PIP problem can be solved in polynomial time, enabling CoEvoLink to scale efficiently to large virus-host datasets. We demonstrate the generalizability of CoEvoLink by applying it to identify putative bat hosts of betacoronaviruses missing in the VIRION database and to predict bacteriophage hosts from metagenomic Hi-C data.

There are several avenues for future research. First, while CoEvoLink currently adopts a maximum parsimony approach, it can be extended to maximum likelihood-based or Bayesian inference, to quantify uncertainty in predicted interactions. Second, while we model virus–host interactions as binary (present/absent) traits, the approach could be generalized to continuous interaction strengths, capturing degrees of compatibility. Third, CoEvoLink can be extended to uncover higher-order ecological interactions [73] of species, such as with Vector-Pathogen-Host complexes [74], which would involve modeling the evolution of interaction tensors on multiple (more than two) phylogenies. Lastly, while we focus on the application of virus-host interactions, the underlying framework applies broadly to inferring any missing interactions between any co-evolving system – for example cancer-immune coevolution [75–77] or plant-insect coevolution [78]. We envision that CoEvoLink will provide a foundation for the future development of such algorithms.

## 7 Data and Code Availability

All data required to reproduce the analyses in this study, along with the complete source code for CoEvoLink, are available in the project GitHub repository:https://github.com/sashittal-group/CoEvoLink. The VIRION dataset used in this study is publicly available on Zenodo at: https://zenodo.org/records/17946436. The betacoronavirus (*β*CoV) dataset is publicly available at: https://github.com/viralemergence/Fresnel/tree/master. The MetaHiC benchmark datasets analyzed in this work are available at: https://github.com/KennthShang/HostPredictionReview/tree/main.

## 8 Author Contributions

P.S. conceived the project, P.S. and M.Z.U.S.C. developed the theory and algorithms. M.Z.U.S.C. implemented the algorithms and performed experimental evaluations. P.S. and T.M.M. supervised the project. All authors wrote the manuscript. All authors read and approved the final manuscript.

## 9 Acknowledgments

This work was partially supported by National Science Foundation award CCF-2412389 (T.M.M.) and the Pandemic Prediction and Prevention Destination Area at Virginia Tech. We further acknowledge support from the Computational Tissue Engineering Interdisciplinary Graduate Education Program (IGEP) at Virginia Tech.

