## Supplementary Materials for "A Cophylogenetic Approach for Virus-Host Interaction Prediction"

**Abstract**

Advances in metagenomics have rapidly expanded viral discovery, revealing vast diversity across Earth's virosphere. Yet most virus–host interactions—i.e., which viruses infect which hosts—remain unrecorded. Identifying these interactions is essential for anticipating zoonotic spillover events and advancing biomedical applications such as bacteriophage therapy. However, the sheer diversity of viruses and hosts makes comprehensive experimental mapping infeasible, motivating the need for computational approaches. Most existing prediction methods rely on supervised learning strategies that use sequence-derived features, such as codon usage bias or $k$-mer frequencies, and do not model the coevolutionary processes that shape virus–host interactions. This limits their ability to generalize and the evolutionary interpretability of their predictions. We introduce CoEvoLink, a framework for predicting virus–host interactions that integrates sequence-based evidence with phylogenetic signal by explicitly modeling the coevolutionary histories of viruses and hosts. CoEvoLink infers likely but unobserved interactions by minimizing the number of evolutionary events required to explain them, yielding the most parsimonious interaction under a coevolutionary model. This formulation generalizes classical maximum parsimony, typically defined on a single phylogeny, by jointly optimizing parsimony across both virus and host phylogenies. Sequence-based information is incorporated by assigning a cost to each potential interaction that reflects its likelihood based on genomic features. By drawing a connection between computing parsimony on interaction matrices and maximum parsimony on phylogenetic networks, we derive a polynomial-time algorithm that balances parsimony with sequence-derived prediction cost. We demonstrate the effectiveness of CoEvoLink on simulated data under diverse coevolutionary models. Applying CoEvoLink, we identified putative bat hosts of betacoronaviruses that have not yet been cataloged in the VIRION database. On a benchmark derived from metagenomic sequencing data, we demonstrate that CoEvoLink improves the performance of existing phage-host prediction tools using cophylogenetic information.

**Code availability:** `https://github.com/sashittal-group/CoEvoLink`

Note: This paper is accepted at RECOMB 2026 (30th Annual International Conference on Research in Computational Molecular Biology).

**Supplementary**

#### Contents


**A Proofs** **S2**

A.1 Proof of Lemma 3.1 and Corollary 3.1 . . . . . S2

A.2 Proof of Theorem 3.1 . . . . . S3

A.3 Proof of Theorem 3.2 . . . . . S4

**B CoEvoLink: Algorithmic details** **S4**

**C Simulation details** **S5**

C.1 Mk Model Simulations . . . . . S5

C.2 Host-Repertoire Model . . . . . S6

**D Supplementary results** **S8**

**E Phylogeny Construction** **S12**

### A Proofs

#### A.1 Proof of Lemma 3.1 and Corollary 3.1

We restate the Lemma and provide a proof.

**Lemma A.1.** *The parsimony score $S(A, T_v, T_h)$ of an interaction matrix $A$ is equal to the hardwired parsimony score of phylogenetic network $N(A, T_v, T_h)$.*

*Proof.* Recall that $N(A, T_v, T_h)$ is a phylogenetic network with $nm$ leaves where each leaf corresponds to a virus-host pair $(i, j)$ and is labeled by entry $a_{i,j}$ of interaction matrix $A$.

Let $\ell$ be a binary vertex labeling on the vertices of $N(A, T_v, T_h)$ that is consistent with the leaf labeling of $N(A, T_v, T_h)$, i.e. $\ell(i, j) = a_{i,j}$. The hardwired parsimony score induced by labeling $\ell$ is the total weight of all edges $((x, y), (x', y'))$, where $\ell(x, y) \neq \ell(x', y')$. The hardwired parsimony score of $N(A, T_v, T_h)$ is the minimum induced hardwired parsimony score over all such possible labelings $\ell$.

It is easy to show that parsimony score $S(A, T_v, T_h)$ of an interaction matrix $A$ is equal to hardwired parsimony score of $N(A, T_v, T_h)$ by observing that there exists a mapping between extensions $\hat{A}$ of $A$ and feasible labelings $\ell$ of $N(A, T_v, T_h)$. Specifically, for every vertex $(x, y)$, we assign $\ell(x, y) = \hat{a}_{x,y}$. Observe that the weight $w_N((x, y), (x', y'))$ of edge $((x, y), (x', y'))$ in $N(A, T_v, T_h)$ is $w_v(x, x')$ when $y = y'$ and $w_h(y, y')$ when $x = x'$. As such, we get the same parsimony score induced by the two labelings,

$$
\begin{aligned}
S(A, T_v, T_h) =& \sum_{j=1}^{m} \sum_{(u,u') \in E(T_v)} w_v(u, u') \mathbf{1}(\hat{a}_{u,j} \neq \hat{a}_{u',j}) + \\
& \sum_{i=1}^{n} \sum_{(u,u') \in E(T_h)} w_h(u, u') \mathbf{1}(\hat{a}_{i,u} \neq \hat{a}_{i,u'}) \\
=& \sum_{j=1}^{m} \sum_{(u,u') \in E(T_v)} w_N((u, j), (u', j)) \mathbf{1}(\ell(u, j) \neq \ell(u', j)) + \\
& \sum_{i=1}^{n} \sum_{(u,u') \in E(T_h)} w_N((i, u), (i, u')) \mathbf{1}(\ell(i, u) \neq \ell(i, u')) \\
=& \sum_{((x,y),(x',y')) \in E(N(A,T_v,T_h))} w_N((x, y), (x', y')) \mathbf{1}(\ell(x, y) \neq \ell(x', y')),
\end{aligned}
$$

where the last term is the hardwired parsimony score of $N(A, T_v, T_h)$ induced by labeling $\ell$. As such, the maximum parsimony score $S(A, T_v, T_h)$ is equal to the hardwired parsimony score of $N(A, T_v, T_h)$.

□

We restate Corollary 3.1 here and provide a proof.

**Corollary A.1.** *The parsimony score $S(A, T_v, T_h)$ of an interaction matrix $A$ with $n$ viruses and $m$ hosts can be computed in $O(nm)$ time.*

*Proof.* Note that all reticulation vertices, i.e. vertices with more than one incoming edges, in $N(A, T_v, T_h)$ are leaves where the labeling is fixed. Furthermore, every leaf $(i, j)$ of the network has exactly two incoming edges. As such, we can construct a tree out of the network $N(A, T_v, T_h)$, where each leaf is split into two nodes, each with one of the incoming edges. The hardwired parsimony score of $N(A, T_v, T_h)$ can then be obtained by applying Sankoff's algorithm on the resulting tree. For completeness, we provide the dynamic program here. Let $f[(x, y), s]$ be the minimum parsimony score obtained in the sub-network in $N(A, T_v, T_h)$ rooted at vertex $(x, y)$ that can be attained if $(x, y)$ is labeled by state $s \in \{0, 1\}$. The following recurrence defines $f[(x, y), s]$.

$$f[(x,y),s] = \begin{cases} 0, & \text{if } x \in L(T_v), y \in L(T_h), \quad a_{x,y} = s, \\ \infty, & \text{if } x \in L(T_v), y \in L(T_h), \quad a_{x,y} \neq s, \\ \displaystyle\sum_{(x',y') \in \delta_N((x,y))} \min_{t \in \{0,1\}} \{\mathbf{1}(s \neq t) + f[(x',y'),t]\}, & \text{otherwise.} \end{cases}$$

The parsimony score $S(A, T_v, T_h)$ of the interaction matrix is given by $\min_{s \in \{0,1\}} f[r(N(A, T_v, T_h)), s]$. □

#### A.2 Proof of Theorem 3.1

We restate the theorem here for completeness and then provide a proof.

**Theorem A.1.** *The Balanced Parsimonious Interaction Prediction Problem can be solved in $O(n^2m^2)$ time, where $n$ is the number of viruses and $m$ is the number of hosts.*

*Proof.* We show this by reducing the BALANCED PIP problem to the minimum $s$-$t$ cut problem on a graph with $O(nm)$ vertices and edges. Since minimum $s$-$t$ cut problem on a graph $G = (V, E)$ can be solved in $O(VE)$ time, this completes the proof.

We start by constructing a graph $G(A', T_v, T_h)$ for a given observed interaction matrix $A'$. The construction is described in the main text, but we re-iterate it here for completeness. $G(A', T_v, T_h)$ is obtained by modifying phylogenetic network $N(A', T_v, T_h)$ (see Section A.1) with the following four steps. First, the root node and orientations of edges of the network are removed. Second, all edge weights are scaled by $\lambda$. Third, source vertex $s$ is added and connected to every vertex $(i, j)$ where $a'_{i,j} = 0$ with edge weight $(1 - \lambda)c_{i,j}$. Fourth, target vertex $t$ is added and connected to every vertex $(i, j)$ where $a'_{i,j} = 1$ with edge weight $\infty$.

We will show that size of the minimum $s$-$t$ cut of $G(A', T_v, T_h)$ is equal to minimum possible $\lambda S(A, T_v, T_h) + (1 - \lambda)c(A, A')$, where $A$ is obtained by performing $0 \to 1$ flips in $A'$.

Consider the minimum $s$-$t$ cut $E'$ in $G(A', T_v, T_h)$, which by definition breaks the graph into at least two components, one of which has $s$ and another of which as $t$. We obtain interaction matrix $A$ and its extension $\hat{A}$ as follows. $A$ is obtained by setting $a_{i,j} = 0$ if $(i, j)$ belongs to the component with vertex $s$, and $a_{i,j} = 1$ if $(i, j)$ belongs to any other component. Similarly, we assign $\hat{a}_{x,j} = 0$ if $(x, j)$ belongs to component with vertex $s$ and $\hat{a}_{x,j} = 1$ otherwise, and $\hat{a}_{i,y} = 0$ if $(i, y)$ belongs to component with vertex $s$ and $\hat{a}_{i,y} = 1$ otherwise. Now, for every edge $(x, x')$ of viral phylogeny $T_v$ where $\hat{a}_{x,j} \neq \hat{a}_{x',j}$ for some host $j$ must be part of the cut $E'$ (since corresponding vertices $(x, j)$ and $(x', j)$ must belong to separate components). Similarly, every edge $(y, y')$ of host phylogeny $T_h$ where $\hat{a}_{i,y} \neq \hat{a}_{i,y'}$ for some virus $i$ must be part of the cut $E'$ (since corresponding vertices $(i, y)$ and $(i, y')$ must belong to separate components). Therefore, the edges corresponding to parsimony score are a subset of $E'$. For every edge where $a'_{i,j} = 0$ and $a_{i,j} = 1$, the set $(s, (i, j))$ must be part of cut $E'$ (since $(i, j)$ is not part of the component with vertex $s$). Therefore, the edges corresponding to prediction (i.e. $0 \to 1$ flips) must also be a subset of $E'$. Since the parsimony edges and prediction edges are disjoint, the total weight of these edges must be less than the total size of the cut. Therefore, $\lambda S(A, T_v, T_h) + (1 - \lambda)c(A, A')$ is less than or equal to the size of the minimum cut of $G(A', T_v, T_h)$.

Consider interaction matrix $A$ that minimizes $\lambda S(A, T_v, T_h) + (1 - \lambda)c(A, A')$ and its most parsimonious extension $\hat{A}$. We use this a cut $E'$ in $G(A', T_v, T_h)$ as follows. If $a_{i,j} = 1$ and $a'_{i,j} = 0$, we include these edges in the cut. It is easy to see that the total cost of these cuts is $c(A, A')$. We also include the edges $((x, y), (x', y'))$ in $G(A', T_v, T_h)$ as part of the cut if $\hat{a}_{(x,y)} \neq \hat{a}_{(x',y')}$. Every path from $s$ to $t$ in $G(A', T_v, T_h)$ must pass through one of the edges of this cut. Therefore, $E'$ is an $s$-$t$ cut. Moreover, the total weight of this cut is equal to $\lambda S(A, T_v, T_h) + (1-\lambda)c(A, A')$. Therefore, minimum $s$-$t$ cut in $G(A', T_v, T_h)$ is less than or equal to the minimum $\lambda S(A, T_v, T_h) + (1 - \lambda)c(A, A')$. This concludes the proof. □

##### A.3 Proof of Theorem 3.2

We restate the theorem here and then provide a proof.

**Theorem A.2.** *The* BOUNDED PIP *problem is NP-hard, even for $n = 1$ or $m = 1$.*

*Proof.* We show that BOUNDED PIP problem is NP-hard with a reduction from the knapsack problem. The Knapsack problem is stated as follows: Given $k$ items with weights $w_1, \ldots, w_k$ and values $q_1, \ldots, q_k$, determine which items should be selected so that the total weight of the selected items is less than or equal to limit $\gamma$ and the total value is maximized. Given weights $w_1, \ldots, w_k$, values $q_1, \ldots, q_k$ and limit $\delta$ we construct an instance of BOUNDED PIP problem as follows. Let number of viruses $n = 1$ and number of hosts $m = k + 1$. We build a $1 \times (k + 1)$ observed interaction matrix $A'$ with $a'_{1,i} = 0$ for all $i$, $1 \times (k + 1)$ cost matrix $C$ where $c_{1,i} = w_i$ for all $i \in [k]$ and $c_{1,k+1} = 0$, and the threshold $\delta = \gamma$. The host tree $T_h$ has one root $r$ and $k + 1$ children corresponding to the entries of $A'$. The weight of edge $(r, i)$ is $q_i$ for $i \in [k]$ and $\sum_i q_i + \epsilon$ for edge $(r, k + 1)$. We show that the knapsack problem has a solution with total value at least $\alpha$ if and only if the constructed BOUNDED PIP problem has a solution parsimony score at most $\sum_i q_i - \alpha$.

($\Leftarrow$) Let $z_i$ be a binary variable that indicates if an item was selected in a solution of the Knapsack problem (0 indicates not selected and 1 indicates selected). We construct a solution of BOUNDED PIP problem by flipping $(1, i)$ entries from $0 \to 1$ if $z_i = 1$. The total cost of all these flips is $\sum_i w_i \leq \gamma = \delta$. Note that since weight of edge $(r, k + 1)$ is greater than the sum of all other edges, the most parsimonious extension $\hat{A}$ of $A$ will always have the root labeled by 1, i.e. $\hat{a}_{1,r} = 1$. As such, the parsimony score of the resulting interaction matrix $A$ is $\sum_i q_i - \sum_i z_i q_i \leq \sum_i q_i - \alpha$.

($\Rightarrow$) Let $A$ be a solution of the BOUNDED PIP problem and $\hat{A}$ be its most parsimonious extension on $T_h$. Since the parsimony score is at most $\sum_i q_i - \alpha$ which is less than $\sum_i q_i + \epsilon$, we must have $\hat{a}_{1,r} = 1$. We obtain a solution to the knapsack problem as follows. We select items based on the entries of interaction matrix $A$, i.e. we select item $i$ if $a_{1,i} = 1$. The total value of these items at least $\alpha$ and the total weight of these items less than $\delta = \gamma$. This concludes the proof. □

#### B CoEvoLink: Algorithmic details

**Enumerating Pareto optimal interaction matrices.** Solving the Pareto Optimal Interaction Prediction Problem (POIP, Prob. 2.3) involves all interaction matrices that Pareto optimal with respect to $S(A, T_h, T_v)$ and $c(A, A')$. We achieve this by solving the BALANCED PIP Problem (Prob. 2.1) for varying values of parameter $\lambda \in [0, 1]$, where each instance is solved in polynomial time using a reduction to the minimum $s$-$t$ cut problem [79]. A key challenge is that several values of $\lambda$ may yield the same solution to the BALANCED PIP (pseudocode in Algorithm 1). We address this by employing the technique described in [80], where a queue of intervals $[\lambda^-, \lambda^+]$ is maintained and we sample $\lambda$ from these intervals only if the BALANCED PIP yields distinct interaction matrices for $\lambda = \lambda^-$ and $\lambda = \lambda^+$ (pseudocode in Algorithm 2).

**Determining the number of missing interactions.** We determine the number of missing interactions by identifying the *elbow* of the Pareto Front that best balances the trade-off between parsimony and prediction cost. To this end, we use *kneedle* [81] a heuristic algorithm that defines the elbow as the the point of maximum curvature on the Pareto front.

**Solving the BOUNDED PIP problem.** For a given threshold $\delta$ on prediction cost, we employ a binary search to find the value of $\lambda$ for which the BALANCED PIP and the BOUNDED PIP problems yield the same solution. Starting with initial bounds $\lambda_{\text{low}} = 0$ and $\lambda_{\text{high}} = 1$, we iteratively solve the BALANCED PIP for $\lambda_{\text{mid}} = (\lambda_{\text{low}} + \lambda_{\text{high}})/2$. If the resulting prediction cost $c(A, A')$ exceeds $\delta$, we set $\lambda_{\text{high}} = \lambda_{\text{mid}}$ and otherwise we set $\lambda_{\text{low}} = \lambda_{\text{mid}}$. We continue this process until prediction cost is equal to $\delta$ within a some tolerance with default of $10^{-2}$ (pseudocode in Algorithm 3).

### C Simulation details

#### C.1 Mk Model Simulations

The Mk model is a generalization of the Jukes–Cantor model for binary traits, where transitions between states occur at rates governed by an instantaneous rate matrix $\mathbf{Q}$ [82]. Specifically, for a two-state character, $\mathbf{Q}$ is defined as:

$$\mathbf{Q} = \begin{pmatrix} -r_{01} & r_{01} \\ r_{10} & -r_{10} \end{pmatrix},$$

where $r_{01}$ is the rate of transition from state 0 to 1, and $r_{10}$ is the rate from 1 to 0. Once this transition rate matrix is specified, the probability distribution of trait states after any time interval $t$ can be obtained from the matrix exponential [57]:

$$\mathbf{P}(t) = e^{\mathbf{Q}t}, \tag{1}$$

where the $(i, j)$-th entry of $\mathbf{P}(t)$ gives the probability of transitioning from state $i$ to state $j$ over time $t$.

For the two-state case, the matrix exponential in Eq. 1 has a closed-form solution:

$$\mathbf{P}(t) = \begin{pmatrix} \frac{r_{10}}{r_{01}+r_{10}} + \frac{r_{01}}{r_{01}+r_{10}} e^{-(r_{01}+r_{10})t} & \frac{r_{01}}{r_{01}+r_{10}}\left(1 - e^{-(r_{01}+r_{10})t}\right) \\ \frac{r_{10}}{r_{01}+r_{10}}\left(1 - e^{-(r_{01}+r_{10})t}\right) & \frac{r_{01}}{r_{01}+r_{10}} + \frac{r_{10}}{r_{01}+r_{10}} e^{-(r_{01}+r_{10})t} \end{pmatrix}.$$

The probability of remaining in the same state can be expressed as the weighted average of the diagonal entries, using the stationary distribution of the Markov process:

$$P_{\text{same}}(t) = \pi_0 P_{00}(t) + \pi_1 P_{11}(t),$$

where the stationary probabilities are

$$\pi_0 = \frac{r_{10}}{r_{01}+r_{10}}, \qquad \pi_1 = \frac{r_{01}}{r_{01}+r_{10}}.$$

Substituting these expressions yields the closed-form solution:

$$P_{\text{same}}(t) = \frac{r_{10}^2 + r_{01}^2 + 2r_{01}r_{10}e^{-(r_{01}+r_{10})t}}{(r_{01}+r_{10})^2}.$$

This quantity captures the overall probability that a binary trait remains unchanged along an evolutionary branch of length $t$. Higher values of $P_{\text{same}}(t)$ indicate lower effective transition rates and greater stability of the trait, reflecting a stronger coevolutionary conservation signal in our model.

To simulate trait evolution, we sampled from $\mathbf{P}(t)$ along the branches of the phylogenetic network and used the resulting states to populate the interaction matrix. The probability $P_{\text{same}}(t)$ served as the edge weight in the network. Thus, higher edge weights correspond to a lower probability of selecting that edge as a cut edge, and therefore a lower chance of changing the state along that branch. In all simulations, a unit prediction cost was assumed for unobserved cells.

In our setting, each entry of the host–virus interaction matrix corresponds to a leaf node that has *two* parents: one from the host phylogeny and one from the virus phylogeny. Because the Mk model and the expression for $\mathbf{P}(t)$ are derived for a single-parent lineage, we extended the transition rule to accommodate two incoming edges. Specifically, if a leaf has parental states $(i, j)$, where $i$ is inherited from the host lineage and $j$ from the virus lineage, then the probability of the leaf being in state $k$ is computed by multiplying the corresponding transition probabilities along both trees. For example,

$$\mathbf{P}((i,j) \to 0) \propto P_{i0}^{(\text{host})}(t_{\text{host}})\, P_{j0}^{(\text{virus})}(t_{\text{virus}}), \qquad \mathbf{P}((i,j) \to 1) \propto P_{i1}^{(\text{host})}(t_{\text{host}})\, P_{j1}^{(\text{virus})}(t_{\text{virus}})$$

with analogous expressions for all four parental combinations $(0,0)$, $(0,1)$, $(1,0)$, and $(1,1)$. These unnormalized probabilities are then normalized to sum to one, yielding a valid distribution for sampling the descendant state. This construction ensures that information from both evolutionary lineages jointly influences the simulated interaction state. We simulated missing data by randomly flipping 2%, 5%, 10%, 20%, and 50% of the 1's in the interaction matrix to 0's.

We explored all combinations of transition rates $r_{01}$ and $r_{10}$ from the set $\{0.1, 0.2, 0.3, 0.4, 0.5, 0.6, 0.7\}$, generating 15 replicate matrices for each pair. Although we present results only for equal-rate pairs and for the 10%, 20%, and 50% missing-data settings, the outcomes for other configurations were similar. Across this entire range of parameters, CoEvoLink consistently recovered missing interactions at comparable levels, demonstrating the robustness of the approach.

##### C.2 Host-Repertoire Model

Braga *et al.* [16] proposed a Bayesian framework for inferring coevolutionary histories between parasites and hosts, modeling the evolution of host repertoires as a continuous-time Markov process. In this model, each parasite taxon has a *host repertoire*, represented by a vector of length $N$ (number of host taxa), where each entry $x_{m,n}$ denotes the interaction state between the $m$-th parasite and the $n$-th host. The state $x_{m,n}$ can be one of three values: 0 (nonhost), 1 (potential host), or 2 (actual host). The state space $S$ includes all repertoires with at least one actual host, resulting in $|S| = 3^N - 2^N$ possible configurations.

Host colonization is modeled as a two-step process: gaining the ability to use a host ($0 \to 1$) and then actually using it ($1 \to 2$). Losses occur in reverse: $2 \to 1$ and $1 \to 0$. Transitions are governed by an instantaneous rate matrix $Q$, where the rate of change from repertoire $y$ to $z$ by altering host $a$ is given by:

$$q_{y,z}^{(a)} = \begin{cases} \mu\lambda_{10} & \text{if potential host loss } (y_a = 1,\, z_a = 0) \\ \mu\lambda_{01}\eta_1(y,a,\beta) & \text{if potential host gain } (y_a = 0,\, z_a = 1) \\ \mu\lambda_{21} & \text{if actual host loss } (y_a = 2,\, z_a = 1) \\ \mu\lambda_{12}\eta_2(y,a,\beta) & \text{if actual host gain } (y_a = 1,\, z_a = 2) \\ 0 & \text{if } |y_a - z_a| > 1 \text{ or repertoires differ in more than one host or } y \text{ has no actual host,} \end{cases}$$

where $\mu$ is the maximum rate of repertoire evolution (rescaling all rates such that $0 \le \mu\lambda_{ij} \le \mu$), and $\lambda_{ij}$ are the base rates for transitions between states $i$ and $j$ (with $0 \le \lambda_{ij} \le 1$). The phylogenetic-distance rate modifiers $\eta_1$ and $\eta_2$ incorporate the effect of host relatedness:

$$\eta(y,a,\beta) = e^{-\beta d/\bar{d}},$$

where $d$ is the average pairwise phylogenetic distance between the new host $a$ and the hosts in $y$ (potential and actual for $\eta_1$, actual only for $\eta_2$), $\bar{d}$ is the average pairwise distance across all hosts, and $\beta \ge 0$ controls the strength of the phylogenetic effect. When $\beta = 0$, host gains are independent of phylogeny; higher $\beta$ favors gains of closely related hosts, increasing phylogenetic conservatism.

In our simulation study, we generated 15 interaction matrix for each combination of $\mu \in \{0.1, 0.5, 1.0\}$ and $\beta \in \{0, 1, 4\}$, using the virus tree and host tree used in the Mk model simulations. The four base transition rates among host-use states ($\lambda_{01}$, $\lambda_{10}$, $\lambda_{12}$, and $\lambda_{21}$) were drawn from a symmetric Dirichlet distribution with concentration parameters $(1, 1, 1, 1)$, i.e., $\boldsymbol{\lambda} \sim$ Dirichlet$(1, 1, 1, 1)$. These simulations produce varying levels of phylogenetic conservatism: low $\mu$ yields similar repertoires among related parasites, while high $\beta$ clusters interactions among related hosts.

To align CoEvoLink with the three-state model used in Braga et al. [16], we adapted their transition rate definitions to our two-state framework (0: no interaction, 2: observed interaction). Following their parameterization, we defined the forward transition rate $r_{01}$ and backward rate $r_{10}$ as:

$$r_{01} = (\lambda_{01} + \lambda_{12})e^{\mu-\beta}, \tag{2}$$

$$r_{10} = (\lambda_{10} + \lambda_{21})e^{\mu}, \tag{3}$$

where $\mu$ scales the overall evolutionary rate, and $\beta$ modulates phylogenetic conservatism in the host tree-higher $\beta$ increases conservatism and reduces transition rates. The parameters $\lambda_{01}, \lambda_{12}, \lambda_{10}, \lambda_{21}$ are switching rates extracted from Braga's simulation settings.

Note that $\beta$ is applied only to $r_{01}$ (gaining interaction) and not to $r_{10}$ (losing interaction). This asymmetry follows Braga et al., who modeled phylogenetic signal in the *acquisition* of host-virus associations but assumed loss events are less constrained by host evolutionary history. This design reflects biological intuition: gaining a compatible viral interaction may require specific co-adaptations (conserved across related hosts), whereas loss can occur via neutral drift or ecological filtering.

Unlike the Mk model simulations with random 1-to-0 flips, here we treat state 1 as *hidden* (potential but unobserved interactions). We collapse observed interactions (state 2) and unobserved non-interactions (state 0) into the known matrix, then use CoEvoLink to infer missing positives (state 1). This evaluates recovery of *latent* host-virus associations under realistic sampling bias.

We explored all combinations of $\mu$ and $\beta$ from Braga's parameter grid, and the results are presented in Figure S1. Across different settings, CoEvoLink achieved an average precision of 0.65, recall of 0.53, and F1-score of 0.6. Specifically, for lower values of $\beta$, the performance decreased due to the sparsity of the interaction matrix. Even in Braga's inference on host-parasite interactions [16], accuracy was lower when estimating the highest rate of host repertoire evolution and lower $\beta$ values.

This shows the efficacy of CoEvoLink under different simulation models. Unlike the Mk model, where missing data are introduced at random, the recovery of potential virus–host interactions here illustrates how the method can be adapted to different settings while still producing strong results.

### D Supplementary results

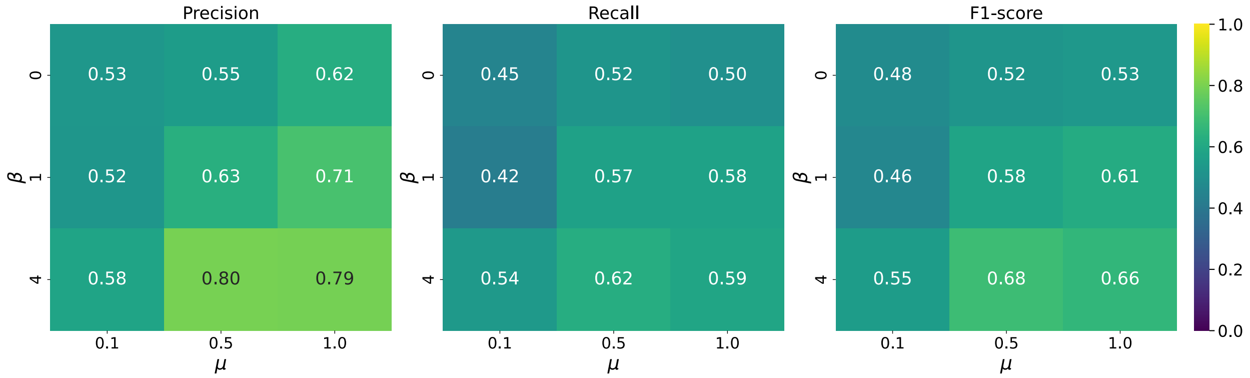


***Fig. S1*** Precision (left), Recall (middle) and F1-score (right) across all combinations of $\mu$(maximum rate of repertoire evolution) and $\beta$(controls the strength of the phylogenetic effect) for the host repertoire model.

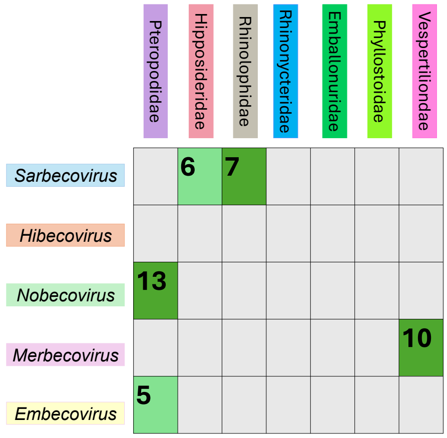


***Fig. S3*** Subgenera-family level interactions predicted by CoEvoLink between betacoronavirus subgenera and bat families. Numbers represent the number of interactions found. Darker colors indicate inference that recapitulates known interactions, and lighter colors indicate inference of new predictions.

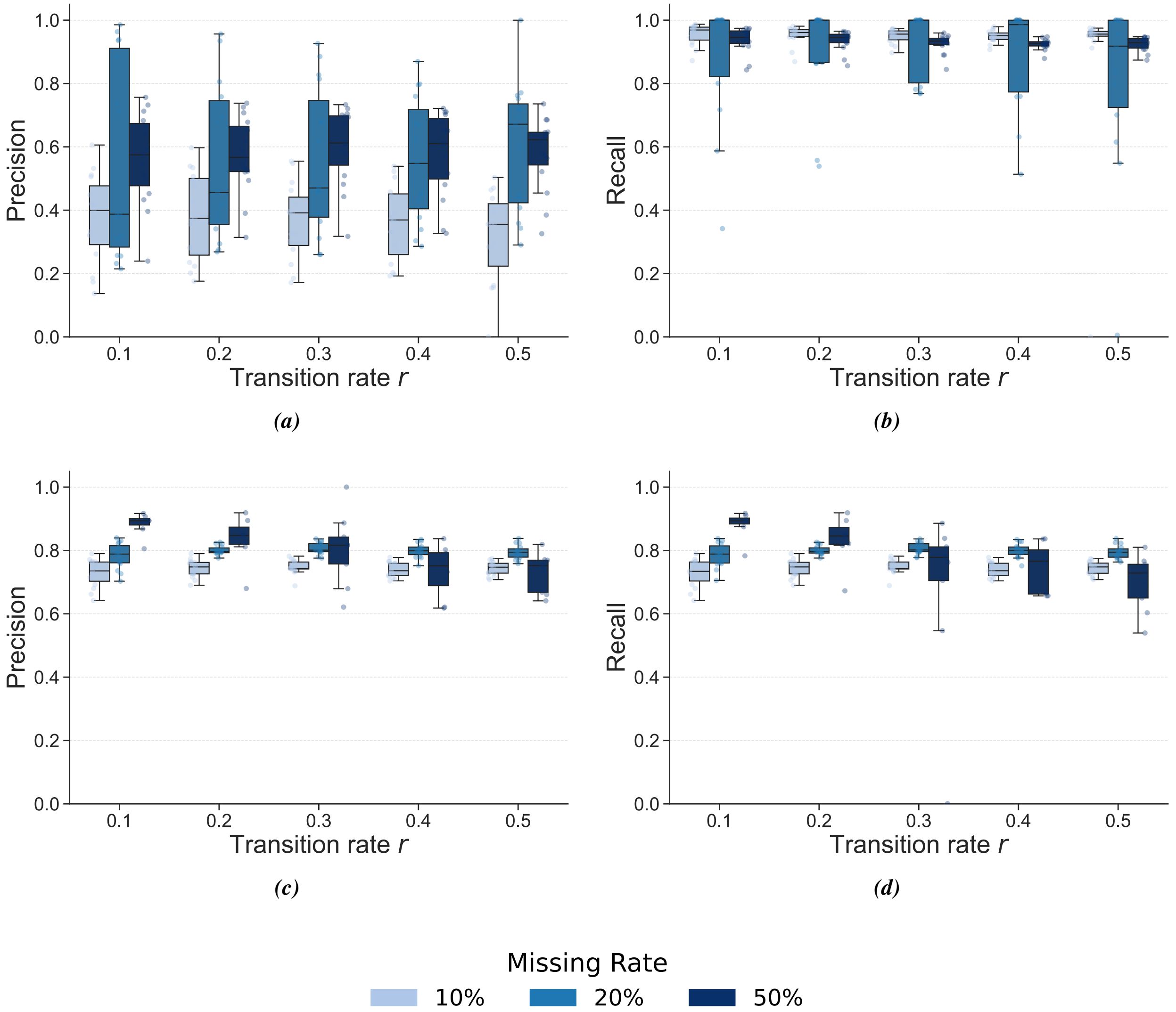


***Fig. S2*** MK model simulation results in (a-b) precision and recall for unsupervised and (c-d) precision and recall for supervised settings across different missing rates.

#### Simulation Results

In Figure S1, precision, recall, and F1 improve with larger values of $\beta$, since lower $\beta$ reduces host conservatism and produces sparser interaction matrices. In Figure S2a (Mk-models), note the drop in precision at low missing rates: in the unsupervised setting, CoEvoLink emphasizes parsimony and therefore predicts more interactions, which lowers precision when the true number of missing entries is small.

#### Performance comparison against genomic-feature-based tools for bacteriophages

From figure S4, it is clear that, across all three datasets (cow fecal, human gut and wastewater samples), integrating CoEvoLink consistently improved or maintained the accuracy of existing host-prediction tools (PHIST, PB-LKS and WIsH) for different *top-k* accuracy.

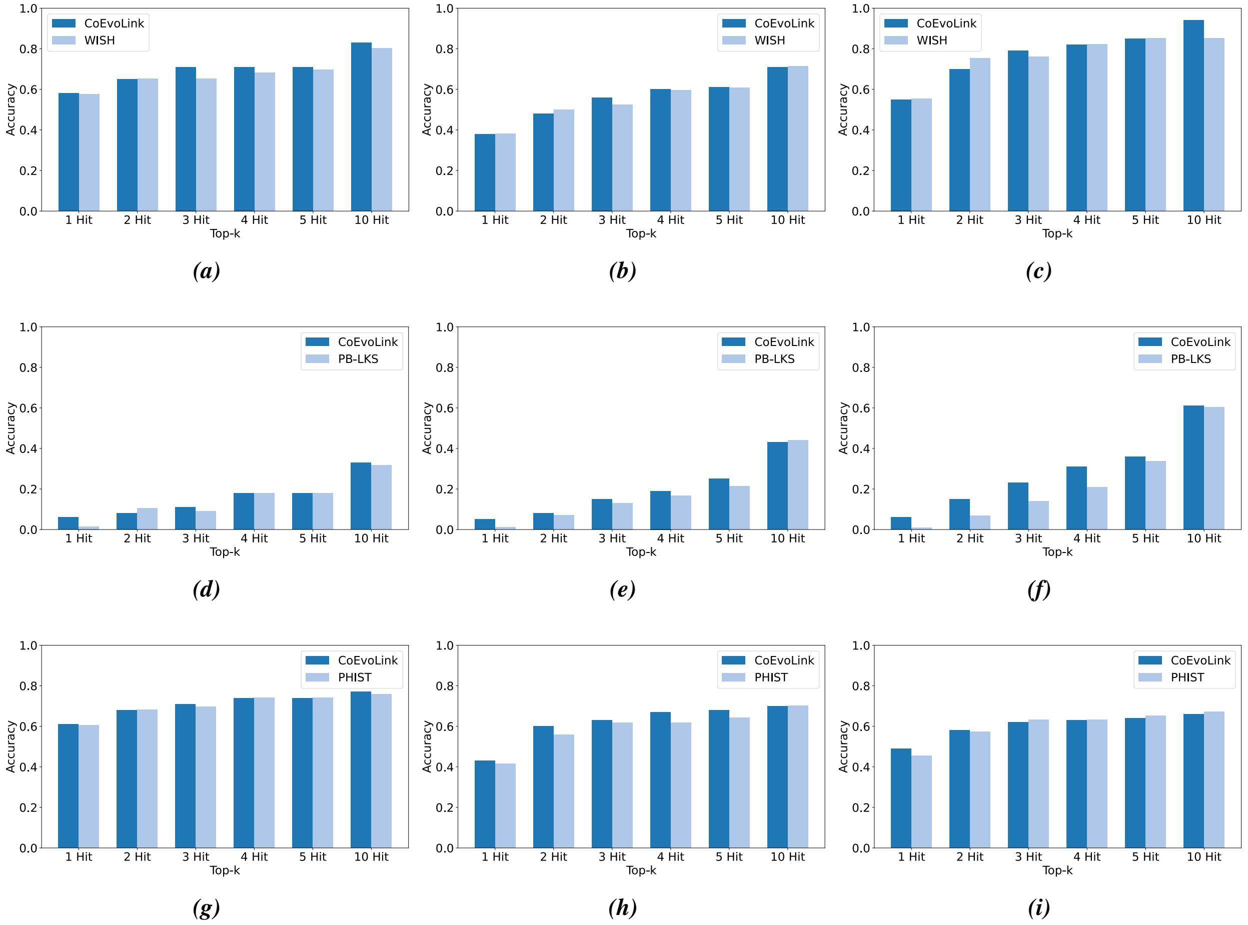


***Fig. S4*** Accuracy of top-k predictions of PB-LKS, PHIST and WIsH with and without CoEvoLink on the interactions from cow fecal (a,d,g), human gut (b,e,h) and wastewater samples (c,f,i).

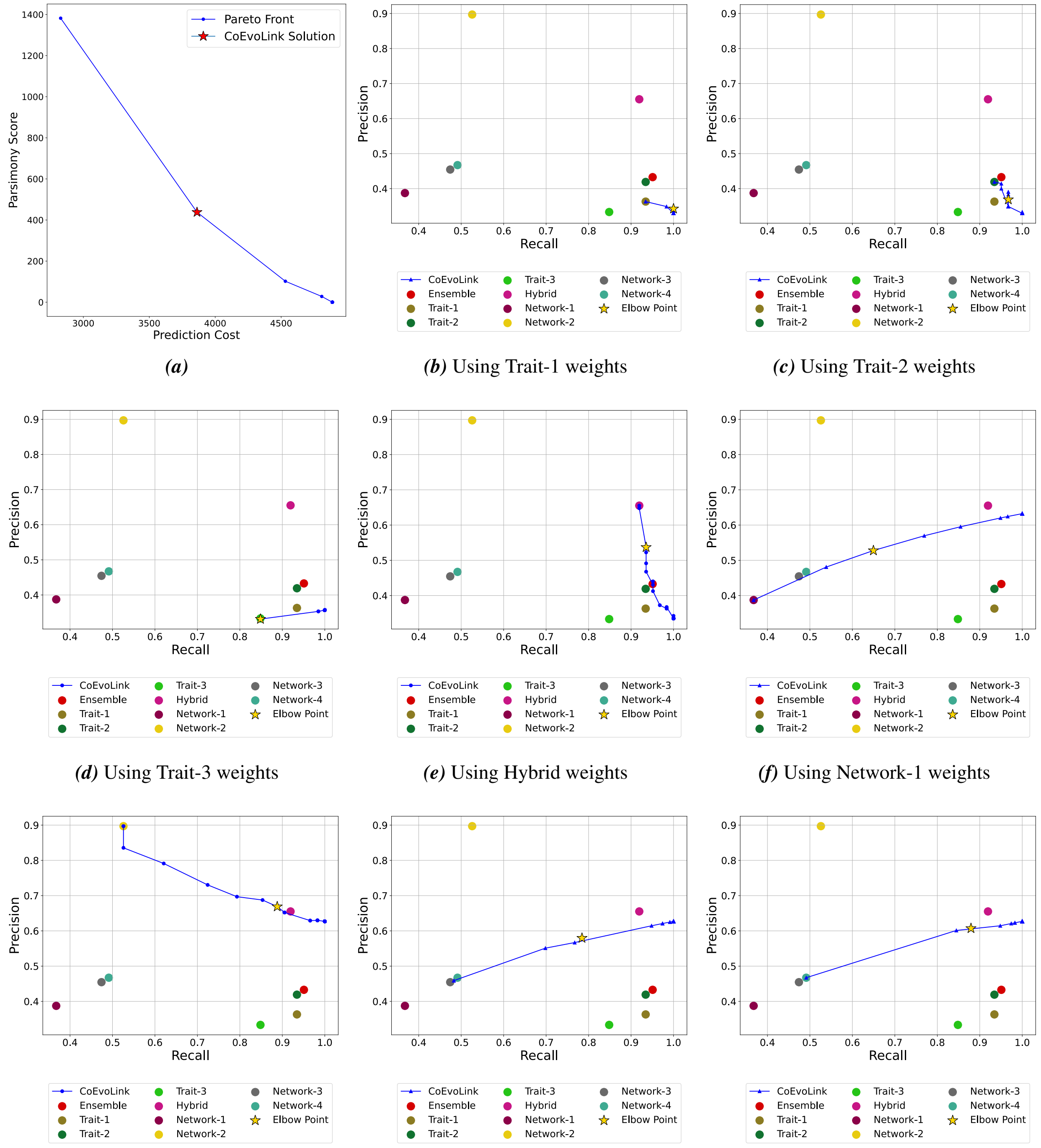


***(a)*** ***(b)*** Using Trait-1 weights ***(c)*** Using Trait-2 weights

***(d)*** Using Trait-3 weights ***(e)*** Using Hybrid weights ***(f)*** Using Network-1 weights

***(g)*** Using Network-2 weights ***(h)*** Using Network-3 weights ***(i)*** Using Network-4 weights

***Fig. S5*** Performance of CoEvoLink across different starting weight configurations. (a) Pareto front showing for using ensemble weights; the selected CoEvoLink solution is marked with a star. (b–i) Precision-recall curves obtained when initializing CoEvoLink with the pretrained weights from each individual model (Trait-1 to Trait-3, Hybrid, Network-1 to Network-4).

#### Performance comparison against eight predictive models

In Figure S5, it is evident that, regardless of the starting point, CoEvoLink consistently achieves substantially higher precision and recall, demonstrating robust improvement over all baseline models. Moreover, for the

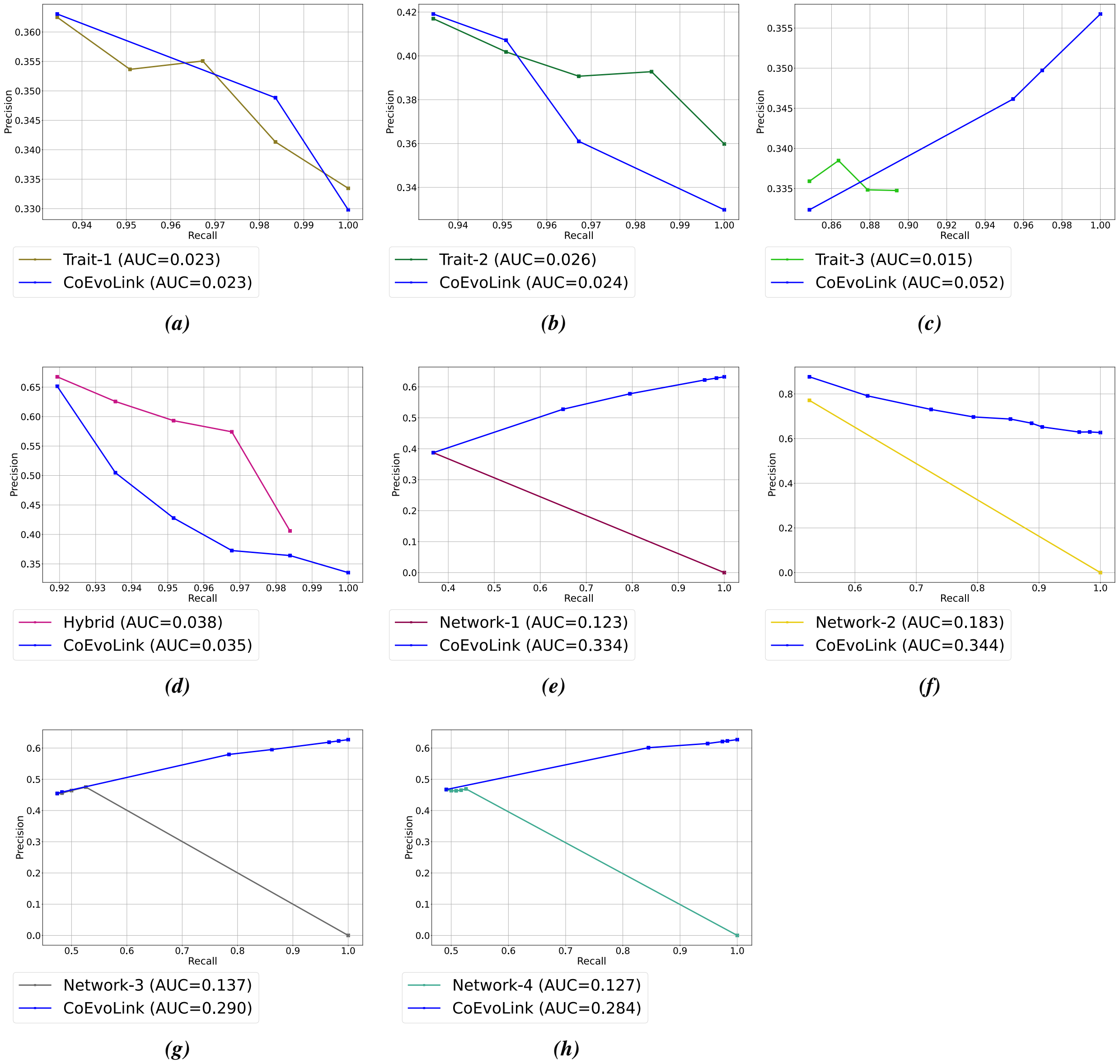


***Fig. S6*** Precision-recall curves comparing the link-prediction performance of CoEvoLink and baseline trait- and network-based methods.

Hybrid and the four network-based models, applying CoEvoLink yields more accurate predictions than the ensemble of all eight models (Supp. Fig. S5(e–i); note that the blue curve lies above the red point representing the ensemble). In Figure S6, apart from Hybrid and Trait-2, CoEvoLink consistently attains higher area under the precision-recall curve than all other approaches.

#### E Phylogeny Construction

To generate phylogenetic trees for all simulation studies, bat–coronavirus analyses, and phage datasets, we implemented a fully automated workflow that retrieves sequence data, filters and preprocesses records, performs multiple sequence alignment, and infers maximum-likelihood phylogenies. All thresholds and parameter settings were kept identical across datasets to ensure methodological consistency.

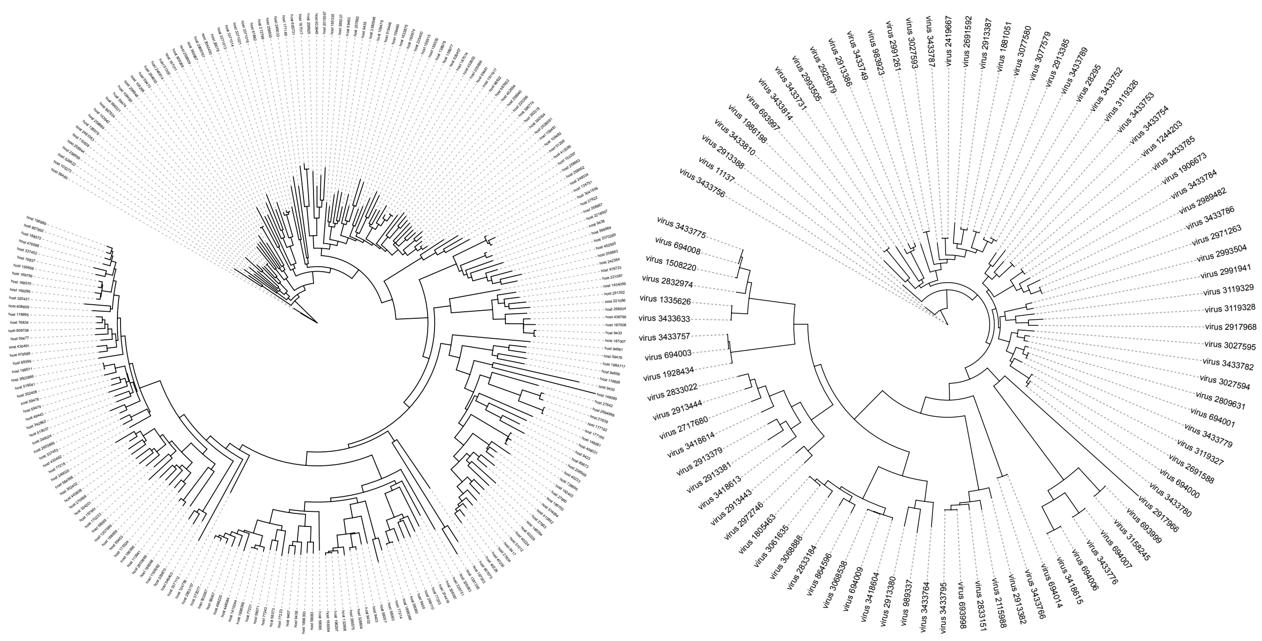

*(a)* *(b)*

***Fig. S7*** (a) Host phylogeny of 233 bat hosts and (b) Virus phylogeny of 94 coronaviruses used in simulation. Virus and host IDs match annotation in VIRION database [33].

#### Data extraction and preprocessing

For analyses involving bats and coronaviruses, we first loaded the complete VIRION database and extracted entries corresponding to host taxa in the order *Chiroptera* and viral taxa in the family *Coronaviridae*. Records were normalized, cleaned, and deduplicated using a composite key of host TaxID, virus TaxID, and accession identifier, ensuring that each host–virus association was represented only once. The resulting filtered table was exported as a metadata file for reproducibility and downstream tracking.

For phage datasets, we began with viral contigs assembled from metagenomic data. Protein-coding sequences were predicted using Prodigal [83] in metagenomic mode (“`-p meta`”). If Prodigal annotations were unavailable or incomplete, contigs were translated in all six reading frames and the longest open reading frame was retained. To preserve alignment quality, contigs shorter than 1000 bp were removed; if a dataset contained no contigs exceeding this threshold, the minimum was relaxed to 500 bp. For phage–host analyses, explicit host genomic sequences were not provided in the benchmark dataset. Therefore, we reconstructed a host phylogeny using taxonomic information from the Genome Taxonomy Database (GTDB). Rather than inferring a sequence-based phylogeny, we generated a taxonomy-based hierarchical tree whose structure reflects GTDB’s standardized bacterial and archaeal taxonomy. Summary statistics, including contig length distributions, were computed to verify dataset quality prior to downstream analysis.

#### Sequence retrieval

Representative viral and host sequences were retrieved programmatically from NCBI using the Entrez API (`Bio.Entrez`), with a fixed delay between queries to comply with NCBI usage guidelines.

For viral taxa, we preferentially downloaded spike glycoprotein sequences from the `protein` database using structured queries of the form:

```
txid<virus_taxid>[Organism:exp] AND (spike glycoprotein OR spike
```

```
protein OR S protein)
```

Only spike sequences of at least 650 amino acids were accepted. If no such sequences were available, we retrieved complete viral genomes from the `nucleotide` database, requiring a minimum length of 20,000 nt.

For host taxa, we queried the `nucleotide` database for complete mitochondrial genomes or common mitochondrial markers (*cytochrome b*, COI (Cytochrome Oxidase I)), using a tiered set of increasingly permissive queries. Retrieved sequences were required to be at least 800 nt in length. All FASTA headers were sanitized to ensure compatibility with alignment and tree-building software.

**Multiple sequence alignment**

Multiple sequence alignments were generated with MAFFT [84] using the `--auto` mode and default settings. Host mitochondrial sequences and viral protein/genome sequences were aligned separately. Before alignment, each dataset was automatically inspected to determine whether sequences were nucleotide or amino acid, ensuring correct handling of viral spike proteins versus nucleotide genomes. The same MAFFT configuration was used across all experiments to maintain uniformity.

**Phylogenetic inference**

Maximum-likelihood phylogenies were inferred using IQ-TREE [56] (version 2 when available), with automatic model selection enabled through ModelFinder ("`-m MFP`"). For each dataset, branch support was assessed using both ultrafast bootstrap replicates and SH-aLRT tests, each performed with 10,000 replicates:

```
-bb 10000 -alrt 10000
```

IQ-TREE was run with automatic thread selection ("`-nt AUTO`"). The resulting Newick-formatted trees, along with associated model reports and logs, were saved to structured output directories. In IQ-TREE, during a phylogenetic analysis with bootstrapping the program typically produces two tree files: `.treefile` – the maximum-likelihood (ML) tree, representing the single best tree found during the search, and `.contree` – the consensus tree, generated from the bootstrap replicates (e.g., a majority-rule consensus tree). This file contains bootstrap support values on its branches. When both `.treefile` and consensus `.contree` files were produced, the consensus tree was retained.

**Reproducibility**

All retrieval thresholds (minimum sequence lengths), Entrez query structures, MAFFT settings, and IQ-TREE parameters were held constant across all datasets and analyses. Jupyter notebooks have been made available on the github repository. This uniform pipeline ensures that any downstream biological signal is not confounded by differences in phylogenetic reconstruction methodology. The resulting host and viral phylogenies served as fixed topologies for all coevolutionary inference and simulation experiments.

| VirusTaxID | Common Name | Virus | VirusGenus | VirusFamily | VirusOrder | VirusClass | VirusNCBIResolved | ICTVRatified | Database |
|---|---|---|---|---|---|---|---|---|---|
| 1805463 | NaN | bat betacoronavirus | betacoronavirus | coronaviridae | nidovirales | pisoniviricetes | True | False | genbank |
| 3061635 | Thomas's horseshoe bat coronavirus | rhinolophus thomasi bat coronavirus | betacoronavirus | coronaviridae | nidovirales | pisoniviricetes | True | False | genbank |
| 3068888 | Thai horseshoe bat sarbecovirus | rhinolophus siamensis bat sarbecovirus | betacoronavirus | coronaviridae | nidovirales | pisoniviricetes | True | False | genbank |
| 2833184 | NaN | sarbecovirus sp. | betacoronavirus | coronaviridae | nidovirales | pisoniviricetes | True | False | genbank |
| 864596 | Blasius's horseshoe bat sarbecovirus | bat coronavirus bm48-31/bgr/2008 | betacoronavirus | coronaviridae | nidovirales | pisoniviricetes | True | False | predict |
| 3068538 | NaN | horseshoe bat sarbecovirus | betacoronavirus | coronaviridae | nidovirales | pisoniviricetes | True | False | genbank |
| 694009 | SARS-CoV-2 | betacoronavirus pandemicum | betacoronavirus | coronaviridae | nidovirales | pisoniviricetes | True | True | eid2; shaw; hp3 |
| 3433764 | Pratt's roundleaf bat coronavirus | betacoronavirus hipposideri | betacoronavirus | coronaviridae | nidovirales | pisoniviricetes | True | True | genbank |
| 989337 | NaN | zaria bat coronavirus | betacoronavirus | coronaviridae | nidovirales | pisoniviricetes | True | False | genbank; predict |
| 2833022 | NaN | nobecovirus sp. | betacoronavirus | coronaviridae | nidovirales | pisoniviricetes | True | False | genbank |
| 2913444 | Madagascar rousette fruit bat nobecovirus | rousettus madagascariensis nobecovirus | betacoronavirus | coronaviridae | nidovirales | pisoniviricetes | True | False | genbank |
| 3418614 | Eidolon bat nobecovirus | betacoronavirus eidoli | betacoronavirus | coronaviridae | nidovirales | pisoniviricetes | True | True | genbank |
| 3418613 | Rousettus bat coronavirus | betacoronavirus cororeum | betacoronavirus | coronaviridae | nidovirales | pisoniviricetes | True | True | genbank |
| 2913443 | Pteropus fruit bat nobecovirus | pteropus rufus nobecovirus | betacoronavirus | coronaviridae | nidovirales | pisoniviricetes | True | False | genbank |
| 1335626 | MERS-CoV | middle east respiratory syndrome-related coronavirus | betacoronavirus | coronaviridae | nidovirales | pisoniviricetes | True | False | predict |
| 2832974 | NaN | merbecovirus sp. | betacoronavirus | coronaviridae | nidovirales | pisoniviricetes | True | False | genbank |
| 3433775 | Pipistrellus bat coronavirus HKU5 | betacoronavirus pipistrelli | betacoronavirus | coronaviridae | nidovirales | pisoniviricetes | True | True | genbank |
| 1928434 | NaN | betacoronavirus sp. | betacoronavirus | coronaviridae | nidovirales | pisoniviricetes | True | False | genbank |
| 3433757 | Human/Cattle coronavirus | betacoronavirus gravedinis | betacoronavirus | coronaviridae | nidovirales | pisoniviricetes | True | True | genbank |
| 3418615 | Rousettus fruit bat coronavirus HKU9 | betacoronavirus rousetti | betacoronavirus | coronaviridae | nidovirales | pisoniviricetes | True | True | genbank |
| 3433776 | Bamboo bat coronavirus (Ty-BatCoV-HKU4) | betacoronavirus tylonycteridis | betacoronavirus | coronaviridae | nidovirales | pisoniviricetes | True | True | genbank |

***Table S1*** Taxonomy for 21 betacoronaviruses in the Virion database. Common names were retrieved through web searches. Some entries appear as *NaN* because the corresponding virus names in Virion are not specific enough to determine common names.

| HostTaxID | Common Name | Host | HostGenus | HostFamily | HostOrder | HostClass | HostNCBIResolved |
|---|---|---|---|---|---|---|---|
| 372077 | Gambian epauletted fruit bat | epomophorus gambianus | epomophorus | pteropodidae | chiroptera | mammalia | True |
| 903567 | Little epauletted fruit bat | epomophorus labiatus | epomophorus | pteropodidae | chiroptera | mammalia | True |
| 77231 | Franquet's epauletted fruit bat | epomops franqueti | epomops | pteropodidae | chiroptera | mammalia | True |
| 58073 | Woermann's bat | megaloglossus woermanni | megaloglossus | pteropodidae | chiroptera | mammalia | True |
| 58063 | Madagascan fruit bat | eidolon dupreanum | eidolon | pteropodidae | chiroptera | mammalia | True |
| 77233 | Niphan's fruit bat | megaerops niphanae | megaerops | pteropodidae | chiroptera | mammalia | True |
| 326081 | Dyak fruit bat | dyacopterus spadiceus | dyacopterus | pteropodidae | chiroptera | mammalia | True |
| 302402 | African roundleaf bat | hipposideros caffer | hipposideros | hipposideridae | chiroptera | mammalia | True |
| 463808 | Sundevall's roundleaf bat | hipposideros ruber | hipposideros | hipposideridae | chiroptera | mammalia | True |
| 58069 | Cantor's roundleaf bat | hipposideros galeritus | hipposideros | hipposideridae | chiroptera | mammalia | True |
| 178895 | Creagh's horseshoe bat | rhinolophus creaghi | rhinolophus | rhinolophidae | chiroptera | mammalia | True |
| 59478 | Geoffroy's horseshoe bat | rhinolophus clivosus | rhinolophus | rhinolophidae | chiroptera | mammalia | True |
| 329870 | Persian trident bat | triaenops persicus | triaenops | rhinonycteridae | chiroptera | mammalia | True |
| 1381355 | Egyptian tomb bat | taphozous perforatus | taphozous | emballonuridae | chiroptera | mammalia | True |
| 27634 | Great fruit-eating bat | artibeus lituratus | artibeus | phyllostomidae | chiroptera | mammalia | True |
| 40228 | Dark fruit-eating bat | artibeus obscurus | artibeus | phyllostomidae | chiroptera | mammalia | True |
| 40233 | Seba's short-tailed bat | carollia perspicillata | carollia | phyllostomidae | chiroptera | mammalia | True |
| 51300 | Noctule bat | nyctalus noctula | nyctalus | vespertilionidae | chiroptera | mammalia | True |
| 246814 | Soprano pipistrelle | pipistrellus pygmaeus | pipistrellus | vespertilionidae | chiroptera | mammalia | True |
| 109485 | Savi's pipistrelle | hypsugo savii | hypsugo | vespertilionidae | chiroptera | mammalia | True |
| 360967 | Great evening bat | ia io | ia | vespertilionidae | chiroptera | mammalia | True |
| 98922 | Daubenton's bat | myotis daubentonii | myotis | vespertilionidae | chiroptera | mammalia | True |
| 249034 | White-bellied yellow bat | scotophilus leucogaster | scotophilus | vespertilionidae | chiroptera | mammalia | True |
| 258952 | African yellow bat | scotophilus dinganii | scotophilus | vespertilionidae | chiroptera | mammalia | True |

***Table S2*** Taxonomy for 24 bat species that are not recorded as betacoronavirus hosts in the Virion database. Common names were retrieved through web searches.

**Algorithm 1** SolveBalancedPIP($\lambda$)

**Input:** Host trees $T_h$, parasite trees $T_v$, precomputed weight matrices $\{W_h\}$ and $\{W_v\}$, prediction cost matrix $C$, current interaction matrix $A'$, and parameter $\lambda \in [0, 1]$.
**Output:** Updated interaction matrix $A$, set of flipped cells, and cut value.

Initialize directed graph $G = (V, E)$ with source $s$ and sink $t$
$H \leftarrow$ list of hosts; $P \leftarrow$ list of parasites
**for** each parasite $p \in P$ with corresponding host tree $T_h^p$ **do**
    $(W, \text{nodes}) \leftarrow W_h^p$
    **for** each edge $(u, v)$ in $T_h^p$ **do**
        $w \leftarrow \lambda \cdot W[u, v]$
        **if** $v$ is a leaf **then**
            node $\leftarrow$ `CELL`_$p$_$v$    ▷ Attach an id to mark as node corresponding to a cell in the matrix
        **else**
            node $\leftarrow v$
        **end if**
        Add bidirectional edges $(u, \text{node})$ and $(\text{node}, u)$ with capacity $w$
    **end for**
**end for**
**for** each host $h \in H$ with corresponding parasite tree $T_v^h$ **do**
    $(W, \text{nodes}) \leftarrow W_v^h$
    **for** each edge $(u, v)$ in $T_v^h$ **do**
        $w \leftarrow \lambda \cdot W[u, v]$
        **if** $v$ is a leaf **then**
            node $\leftarrow$ `CELL`_$v$_$h$
        **else**
            node $\leftarrow v$
        **end if**
        Add bidirectional edges $(u, \text{node})$ and $(\text{node}, u)$ with capacity $w$
    **end for**
**end for**
▷ Attach source/sink based on flip costs and current cell states
**for** each cell $(p, h)$ **do**
    node $\leftarrow$ `CELL`_$p$_$h$
    **if** $A'[p, h] = 0$ **then**
        Add edge $(s, \text{node})$ and $(\text{node}, s)$ with capacity $(1 - \lambda) \cdot C[p, h]$
    **else**
        Add edges $(\text{node}, t)$ and $(t, \text{node})$ with capacity $\infty$
    **end if**
**end for**
Compute minimum $s$-$t$ cut on $G$ to obtain partitions $(R, \bar{R})$
Update cell states: $A[p, h] \leftarrow 0$ if `CELL`_$p$_$h$ $\in R$, else 1
Record flips where $A'[p, h] \neq A[p, h]$
Compute root state for each parasite tree: $r_h \leftarrow 0$ if root $\in R$, else 1
**return** $(A, \text{flips}, \text{cut value}, \{r_h\})$

**Algorithm 2** Enumerate Pareto Optimal Interaction Matrices (ENUMERATEPOIP)

**Input:** Host tree $T_h$, parasite tree $T_v$, observed matrix $A'$
**Output:** Set of Pareto-optimal interaction matrices $\mathcal{A}^*$

Initialize queue $\mathcal{Q} \leftarrow \{\, [0, 1] \,\}$
Initialize $\mathcal{A}^* \leftarrow \emptyset$
**while** $\mathcal{Q}$ is not empty **do**
    Pop interval $[\lambda^-, \lambda^+]$ from $\mathcal{Q}$
    $A^- \leftarrow$ SOLVEBALANCEDPIP$(\lambda^-, T_h, T_v, A')$
    $A^+ \leftarrow$ SOLVEBALANCEDPIP$(\lambda^+, T_h, T_v, A')$
    **if** $A^- \neq A^+$ **then**
        $\lambda^* \leftarrow (\lambda^- + \lambda^+)/2$
        $A^* \leftarrow$ SOLVEBALANCEDPIP$(\lambda^*, T_h, T_v, A')$
        $\mathcal{A}^* \leftarrow \mathcal{A}^* \cup \{A^*\}$
        Push $[\lambda^-, \lambda^*]$ and $[\lambda^*, \lambda^+]$ onto $\mathcal{Q}$
    **end if**
**end while**
**return** $\mathcal{A}^*$

**Algorithm 3** BoundedPIP($\delta$)

**Input:** Host and parasite trees, weight matrices $\{W_h\}, \{W_v\}$, flip cost matrix $C$, observed interaction matrix $A'$, target prediction cost $\delta$, tolerance $\epsilon$, and maximum iterations $K$.
**Output:** Regularization parameter $\lambda^*$ such that $c(A, A') \approx \delta$, and corresponding interaction matrix $A$.

Initialize $\lambda_{\text{low}} \leftarrow 0$, $\lambda_{\text{high}} \leftarrow 1$
Initialize $\lambda^* \leftarrow$ None, $A^* \leftarrow$ None
**for** $k = 1$ to $K$ **do**
    $\lambda_{\text{mid}} \leftarrow (\lambda_{\text{low}} + \lambda_{\text{high}})/2$
    $(A, \text{flips}) \leftarrow$ SOLVEBALANCEDPIP$(\lambda_{\text{mid}})$
    Compute prediction cost $c(A, A') \leftarrow |\text{flips}|$    ▷ or total weighted flip cost if applicable
    **if** $|c(A, A') - \delta| \leq \epsilon$ **then**
        $\lambda^* \leftarrow \lambda_{\text{mid}}$, $A^* \leftarrow A$
        **break**
    **else if** $c(A, A') > \delta + \epsilon$ **then**
        $\lambda_{\text{high}} \leftarrow \lambda_{\text{mid}}$
    **else**
        $\lambda_{\text{low}} \leftarrow \lambda_{\text{mid}}$
    **end if**
    **if** $A^* =$ None or $|c(A, A') - \delta| < |c(A^*, A') - \delta|$ **then**
        $\lambda^* \leftarrow \lambda_{\text{mid}}$, $A^* \leftarrow A$
    **end if**
**end for**
**return** $(\lambda^*, A^*)$

[39] Gustavo Fermin. Host range, host–virus interactions, and virus transmission. *Viruses*, page 101, 2018.

[45] David L Swofford. Phylogenetic analysis using parsimony. 1998.

[67] Maximilian O Press, Andrew H Wiser, Zev N Kronenberg, Kyle W Langford, Migun Shakya, Chien-Chi Lo, Kathryn A Mueller, Shawn T Sullivan, Patrick SG Chain, and Ivan Liachko. Hi-c deconvolution of a human gut microbiome yields high-quality draft genomes and reveals plasmid-genome interactions. *biorxiv*, page 198713, 2017.

[76] Long Wang, Christo Morison, and Weini Huang. Cancer–immune coevolution dictated by antigenic mutation accumulation. *Elife*, 14:RP103970, 2025.
